# Fluorescence based microviscosity mapping in membraneless organelles

**DOI:** 10.64898/2026.01.05.697638

**Authors:** Avantika Kapoor, Pascal Didier, Carla Faivre, Chloé Frey, Pierre Hener, Tania Stefan, Thiebault Lequeu, Eleonore Réal, Andrey Klymchenko, Nicolas Anton, Mayeul Collot, Halina Anton

**Affiliations:** Laboratory of Bioimaging and Pathologies, UMR 7021, CNRS/Université de Strasbourg, 74 route du Rhin, 67401 Illkirch-Graffenstaden, France; Chemistry of Photoresponsive Systems, Laboratoire de Chémo-Biologie Synthétique et Thérapeutique (CBST) UMR 7199, CNRS, Université de Strasbourg, 67400 Illkirch, France; In vitro imaging platform – centre de Neurochimie, UAR 3156, CNRS/ Université de Strasbourg, 8 allée du Général Rouvilloisn 67000 Strasbourg, France; INSERM (French National Institute of Health and Medical Research), UMR 1260, Laboratoire de génétique médicale U1112, Université de Strasbourg, 67000 Strasbourg, France; INSERM (French National Institute of Health and Medical Research), UMR 1260, Regenerative Nanomedecine (RNM), FMTS, Université de Strasbourg, 67000 Strasbourg, France

## Abstract

Membraneless organelles (MLOs) are cellular biomolecular condensates formed by liquid-liquid phase separation. Their biological functions are intimately linked to their material properties including viscosity. Condensate viscosity is determined by the size, shape, concentration and molecular interactions between MLOs components. It impacts the diffusion of MLOs constituents and the selective permeability of the condensate, thereby regulating the rate of biochemical reactions. Viscosity modifications associated with liquid-to-gel transition of the condensates are related to pathologies.

Current experimental approaches for characterizing the material properties of cellular condensates remain limited. In this study, we report the use of BODIPY-based molecular rotors, in combination with fluorescence lifetime imaging microscopy (FLIM), to monitor the microviscosity of cellular MLOs directly in living cells. The fluorescence lifetime of BODIPY derivatives increases with the viscosity of their microenvironment, enabling quantitative assessment of microviscosity within condensates. HaloTag technology was employed to specifically label MLO components.

Our findings reveal that the nucleolus exhibits higher viscosity than the surrounding nucleoplasm and that microviscosity varies across nucleolar sub-compartments. Furthermore, nucleolar reorganization induced by inhibition of rRNA synthesis results in a measurable increase in microviscosity. Finally, we demonstrate that the microviscosity of stress granules is lower than that of the nucleolus.

Overall, presented results demonstrate the strong potential of the BODIPY based molecular rotors as a versatile and powerful tools for probing the material properties of cellular MLOs.

## Introduction

Liquid-liquid phase separation (LLPS) has recently emerged as a fundamental mechanism underlying cellular organization, driving the formation of biomolecular condensates, lipid membrane domains, and selective barriers.^1^ LLPS is a thermodynamically driven, reversible process in which components of a homogeneous solution demix to form dense liquid condensates dispersed within a dilute phase. In the cells LLPS gives rise to organelles, called membraneless organelles (MLOs), such as stress granules, p-bodies, Cajal bodies or nucleolus.^2,3^

MLOs are complex, heterogeneous condensates that behave like fluids: they fuse, coalesce and drip, which are behaviors governed by their physicochemical characteristics (*i.e*. surface tension, viscosity, etc.). Due to their LLPS nature, MLOs represent transient, dynamic, and open systems whose biological functions are tightly linked to their composition, architecture, and material properties. These parameters are of crucial importance, as the internal environment of condensates must be precisely regulated to fulfill their biological roles.^4^

Viscosity is one of the keys physicochemical properties of molecular condensates. Viscosity represents the resistance of a fluid to flow. On a molecular scale this property is determined by concentration, size, shape, and interactions of the fluid’s components. In MLOs, viscosity is particularly important because it affects internal dynamics and diffusion of their components. Condensate viscosity regulates the interfacial properties responsible for the droplet shape and the selective permeability which in turn impacts the rate id biochemical reactions. Under certain conditions, MLOs may undergo further phase transitions into gel-like or solid states, which are often associated with patholgies.^5^ Therefore, understanding the relationship between biological function and the viscoelastic properties of MLOs remains a major challenge.

Several experimental approaches have been developed to probe condensate rheology at different scales. Passive microrheology, based on tracking fluorescent beads within reconstituted droplets, enables viscosity quantification in controlled environments, while droplet coalescence dynamics provide complementary estimates of interfacial tension and viscosity.^6^ Active micro-rheology using optical tweezers further allows direct measurement of viscoelastic properties by applying oscillatory forces to embedded particles. ^7–10^

Compared to *“in vitro”* systems, viscosity measurements in cellular condensates are more fastidious. In living cells, fluorescence recovery after photobleaching (FRAP) has been widely used to assess component mobility and infer viscosity from diffusion parameters.^11,12^ More recently, optical diffraction tomography, a label-free imaging technique exploiting refractive index contrast, has been applied to quantify condensate density.^13,14^ Together, these approaches have expanded our understanding of condensate material states, yet mapping viscosity at the nano to micro scale within living cells remains challenging.

In this work, we introduce a complementary method for measuring the microviscosity within cellular condensates. We designed and optimized a BODIPY-based molecular rotor compatible with fluorescence lifetime imaging microscopy (FLIM), enabling direct, quantitative readout of local microviscosity. BODIPY dyes are environment-sensitive fluorophores, also called “molecular rotors” whose excited-state relaxation depends on environmental viscosity. In low-viscosity media, energy relaxation occurs via non-radiative intramolecular rotation, whereas in viscous environments, restricted motion leads to increased fluorescence quantum yield and lifetime. BODIPY-based rotors have already demonstrated high sensitivity for probing viscosity in cellular compartments such as the plasma membrane, mitochondria, and endoplasmic reticulum, ^15–19^ as well as in protein aggregates.^20^ However, their potential for investigating cellular MLOs has remained unexplored.

To address this gap, we developed a chemogenetic labeling strategy based on a BODIPY derivative functionalized with a chloroalkane ligand for covalent conjugation to HaloTag fusion proteins.^21^ This approach enables selective labeling of condensate components and direct measurement of local viscosity in living cells. Using this tool, our experiments revealed distinct viscosity distributions within nucleolar sub-compartments and dynamic changes accompanying nucleolar reorganization upon inhibition of RNA transcription. Furthermore, we identified significant viscosity differences between nucleoli and stress granules, reflecting their distinct internal architectures. Together, these findings establish BODIPY-based molecular rotors as a powerful tool for live-cell imaging of microviscosity in biomolecular condensates, offering new insights into the material properties that underlie cellular organization.

## Materials and Methods

### Chemicals and Synthesis

All starting materials were purchased from Sigma-Aldrich, BLD Pharm, or TCI Europe and were used as received unless otherwise stated.

NMR spectra were recorded on a Bruker Avance III 400 MHz spectrometer. Chemical shifts (δ) are reported in ppm relative to residual solvent signals. High-resolution mass spectra were acquired using an Agilent Q-TOF 6520 mass spectrometer. Detailed synthetic protocols and full characterization of all compounds are provided in the Supplementary Materials and Methods information.

### Preparation of Glycerol–Water Mixtures for Rotor Calibration

To calibrate the viscosity sensitivity of the BODIPY-based rotor, a series of glycerol-water mixtures with different viscosities were prepared. Glycerol (Euromedex, France) and ultra-pure MilliQ were mixed with defined mass ratios to achieve final glycerol percentages (w/w) 0%, 35%, 45%, 60%, 67%, 72%, 75%, 77%, 82%, 85%, 90% and 95%. Glycerol mixtures with 10 mM Tris, 150 mM NaCl at pH 7.5 were prepared to assess the influence of protein binding under physiological buffer conditions. The viscosity of each solution was measured at 20°C using a HAAKE MARS rotational rheometer (Thermo Scientific, Waltham, MA, USA) equipped with a 35 mm parallel-plate geometry (P35/Ti/SB) set to a 0.10 mm gap and operated with a solvent trap to minimize evaporation. Approximately 100 µL of sample was loaded onto the lower plate and allowed to equilibrate at 20°C prior to measurement. Steady-shear flow curves were acquired in controlled shear-rate mode, applying shear rates from 0.1 to 1000 s⁻¹ and recording the corresponding shear stress. The apparent viscosity was calculated as η = τ⁄γ̇ (where γ̇ is shear rate, τ the shear stress), and, for such Newtonian mixtures, taken from the shear-rate-independent plateau. Measurements were performed in triplicate.

### Fluorescence Quantum yield measurements

The quantum yield (QY) of the BODIPY acid was measured for G0%, G45%, G67%, G77%, G85% and G95% using the integration sphere (SC-30) module of FS5 fluorometer from Edinburgh Instruments. This module allows to determine the absolute fluorescence quantum yield of the dye. A dye concentration of 500 nM was used in a total sample of 4 ml. The sample was heated overnight at 50°C to allow the dye to mix properly at high viscosity (i.e. G77%, G85% and G95%). The excitation wavelength was set to 430 nm and the emission was recorded from 440 to 700 nm.

### Fluorescence Lifetime Measurements

Time-resolved fluorescence measurements were performed with the time-correlated single-photon counting technique. Excitation pulses at 500 nm were generated by a supercontinuum laser (NKT Photonics SuperK Extreme) with 10 MHz repetition rate. The fluorescence decays were collected at 520 nm using a polarizer set at magic angle and a 16 mm band-pass monochromator (Jobin Yvon). The single-photon events were detected with a micro-channel plate photomultiplier R3809U Hamamatsu, coupled to a pulse pre-amplifier HFAC (Becker-Hickl GmbH) and recorded on a time-correlated single photon counting board SPC-130 (Becker-Hickl GmbH). The measured decays were fitted by using a function corresponding to an exponential decay convolved with a normalized Gaussian curve of standard deviation σ standing for the temporal IRF and a Heavyside function. The fitting function was built in Igor Pro (Wavemetrics). All emission decays were fitted using a weighting that corresponds to the standard deviation of the photon number squared root.

Glycerol-water solutions were incubated at room temperature overnight to ensure homogeneity. A stock solution of 200 µM BODIPY rotors was prepared in DMSO. For each glycerol-water mixture, the BOD-L, BOD-PEG4-L, and BOD-PEG12-L rotors were added to achieve a final concentration of 50 nM.

### Cell Culture

Human Embryonic kidney (HEK293) cell lines and Human bone osteosarcoma epithelial cell lines (U2OS) were used for live cell imaging experiments. HEK293T cells were grown in Dulbecco’s Modified Eagle Medium (DMEM,Gibco) supplemented with 10% foetal bovine serum (FBS), 100 μg/mL penicillin, streptomycin, 2 mM L-glutamine and 1 mM sodium pyruvate. U2OS cells were grown in modified Mc’Coy media containing 10% FBS, 100 μg/mL penicillin, streptomycin and 1 mM sodium pyruvate. All cell lines were cultured at 37°C in humidified atmosphere containing 5% CO2.

For fluorescence imaging, cells were seeded onto a 35 mm Ibidi ibi-treat or glass bottom dish. HEK293T cells were seeded with density of 1.5 × 10^5^ cells while U20S were seeded with a density of 5 × 10^4^ cells/dish in 2 mL of their respective media. 24 hours post seeding; cells were transfected with plasmid DNA using Jet PEI transfection kit according to supplier’s protocol.

pcDNA-NPM-HaloTag^22^ and pcDNA-Fib-HaloTag were cloned from eGFP-NPM1 (Addgene n°:17578) kindly provided by Dr. Wang ^23^ and eGFP-Fib (Addgene n°:26673) provided by Dr. Chen.^24^ The sequence coding for HaloTag was amplified from pSEMS-Halo7Tag-hFis (111136 Addgene) vector provided by Dr. Karin Busch ^25^. After purification, the PCR products were digested by BamHI/Xho restriction enzymes and inserted in pcDNA3.1 (zeo) vector. The obtained pcDNA-HaloTag was further digested with HindIII/Bam-HI restriction enzymes and ligated with the NPM or Fib inserts amplified from eGFP-NPM1 and eGFP-Fib plasmids respectively. The final pcDNA-NPM-HaloTag was ligated and sequenced for verification.

Fib-SNAP plasmid was provided by Dr. Castano and Dr. Kriz^26^ and HaloTag-G3BP1 by Dr. Brands’ lab.^27^ Plasmids coding for eGFP1 and eGFP3 were provided by Dr. Nalaskowski.^28^ Plasmids coding for free HaloTag, HaloTag-H2B, HaloTag-LifeAct were kindly provided by Dr. Gauthier. ^29^

Inhibition of RNA Pol 1 was performed by incubating the cells during 2 hours in complete medium containing 2µg/mL Actinomycin D. The stress granules formation was induced by addition of 500µM NaAsO_2_ into the complete growth medium during 1 hour. Cellular RNAs were stained by Pyronin Y (1µM, 15minutes, 37°C). Actinomycin D, NaAsO_2_ and Pyronin Y were purchased from Sigma-Aldrich.

Janelia fluor dyes coupled to Halo and Snap ligands were provided by Janelia materials (HHMI Janelia Research Campus).

### Fluorescence lifetime imaging

Approximately 24 hours post transfection, the growth media was replaced with Opti-MEM (Gibco), and cells were incubated with 200 nM of BODIPY rotor dye for 15-20 minutes at 37°C. Excess dye was removed by washing with Opti-MEM.

FLIM measurements were performed on a Leica Stellaris equipped with a Falcon module and on a homemade two-photon scanning FLIM microscope.

Commercial inverted confocal laser scanning microscope (STELLARIS 8, Leica Microsystems, Nanterre, France) was equipped with a fully fast integrated FLIM module, the so-called FAst Lifetime CONtrast (FALCON, Leica Microsystems, Nanterre, France) and a white light laser (WLL2 440−790 nm). Acquisitions were performed through a 512×512 image format, a scan speed at 400 Hz and a 63X (NA 1.4) oil immersion objective. Bod-4PEG-L imaging was performed with 488 nm excitation (WLL, 8% power). In a photon counting mode, hybrid PMT HyD-X detector was used to detect fluorescence emission from 505 to 750 nm. FLIM images were acquired with accumulation of 8 lines and 3 frames repetitions to detect ∼1000 photons in each pixel showing Bod-PEG4-L labelled structures. A minimum of 100 photons was set to pixels represented in the phasor plot.

The homemade multiphoton scanning microscope is based on an inverted microscope (IX83, Olympus) with a 60X 1.2 NA water immersion objective operating in the descanned fluorescence collection mode.^30^ The BODIPY derivatives were excited at 780 nm using a femtosecond laser (Insight DeepSee, Spectra Physics). Fluorescence photons were collected at a dwell time of 4 µs/pixel using a short-pass filter with a cutoff wavelength of 720 nm (Semrock, FF01-720/SP-25). The fluorescence was directed to a fiber-coupled APD (SPCM-AQR-14-FC, Perkin Elmer), which was connected to a time-correlated single photon counting module (SPC830, Becker & Hickl). The measurement was controlled by SPCM ver 9.83 (Becker & Hickl). The time-resolved fluorescence decay at each pixel was analyzed using a commercial FLIM analysis software package Becker and Hickl SPCImage. The decay curves were fitted using a biexponential model, convolved with the instrument response function (IRF), as:

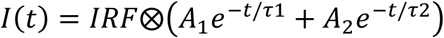

where I(t) represent the intensity decay, IRF denotes the instrument response function of the system. *A*_1_ and *A*_2_ are the amplitudes of the two decay components with *τ*_1_ and *τ2* fluorescence lifetimes. The average fluorescence lifetime τ was then calculated as:

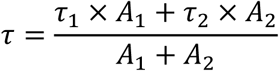

Pixel-wise fitting was carried out to generate fluorescence lifetime maps, and regions of interest (ROIs) were selected to extract quantitative data for comparison across experimental conditions.

### Phasor analysis

Phasor analysis is a fit-free technique in which the fluorescence decay from each pixel is transformed into a point in two-dimensional (2-D) phasor space.^31,32^ If P(i,j) represents a pixel in the FLIM image with coordinates (i,j) and Ii,j(t) is the fluorescence intensity decay at that pixel, the corresponding coordinates in the phasor plot (g,s) for time-domain measurements are given as:

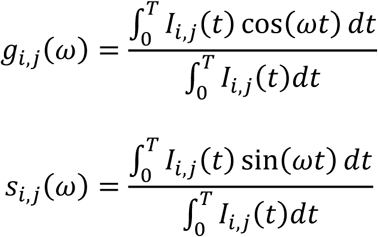

where *ω = 2πf* and *f = 1/T* is the laser repetition rate. Phasor analysis provides a visual distribution of the molecular species in an image by clustering pixels with similar lifetimes.

### Recombinant Protein Production

Recombinant HaloTag protein with a N-terminal 6×-His tag was purified using the *E. coli* expression system, BL21(DE3) cells. Competent bacteria were transformed with pET28-Halo plasmid, and protein expression was induced by adding 500µM IPTG. After 4h of culturing the cells were centrifuged and the pellet was conserved at –80°C. For the purification the cells were lysed by ultrasonication: 2sec ON, 2sec OFF (1 min/mg of dry pellet), 17W/wave, in Lysis buffer (20 mM Tris-HCl, pH 7.5, 150 mM NaCl, 10 mM imidazole, mM β-mercaptoethanol, 1mM PMSF and a protease inhibitor mixture (Roche Diagnostics), then centrifuged at 20000 g, 4°C during 45 minutes. The supernatant was filtered with 0,22 µm low binding filters and loaded on Ni-NTA agarose column (Qiagen) beforehand equilibrated with Equilibration Buffer (20 mM Tris-HCl, pH 7.5, 500 mM NaCl, 15mM imidazole). HaloTag protein was eluted with Ni-Elution buffer (20 mM Tris, pH 7.5, 500 mM NaCl and 1000 mM imidazole). Elution from Ni-NTA was concentrated to 2mL (Amicon Ultra 4, 10K, Millipore) and loaded into size exclusion chromatography column (SEC) beforehand equilibrated with SEC Equilibration Buffer (10 mM Tris, pH 7.5,150mM NaCl, 2mM DTT). Fractions containing Halo protein were pooled, concentrated to ∼ 50 μM and aliquots were flash frozen in liquid nitrogen and stored at −80°C.

## Results

### BODIPY derived molecular rotors are sensitive to viscosity

BODIPY-based molecular rotors display absorption and emission maxima centered at 480 nm and 520 nm respectively. The de-excitation pathway of these molecules depends on conformational flexibility and the rotational freedom of their phenyl moiety (Figure 1A, B). In viscous environments, where molecular mobility is restricted, radiative relaxation is favored, resulting in an increase in both fluorescence quantum yield and lifetime. Hence, we first characterized the viscosity response with water/glycerol mixture of varying viscosity by measuring the fluorescence quantum yield (Φ_F_) of the BODIPY-COOH (BOD-COOH) derivative. The viscosities (η) of the mixtures were systematically determined using an oscillatory rheometer. The relationship between τ, Φ_F_, and η follows the Förster–Hoffmann model as previously described:

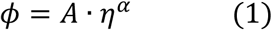

where *A* is a constant and *α* reflects the sensitivity of the molecular rotor to viscosity. The viscosity values ranged from 1 mPa.s for pure water to 506 mPa.s for 95% glycerol at 20°C. As shown in figures 1C and 1E, the fluorescence quantum yield increased with environmental viscosity, following the Förster–Hoffmann relationship with *α* values of 0.71 ± 0.02. We next measured the time resolved fluorescence decays of the same water/glycerol mixtures. In line with the quantum yield measurements, upon increase of the viscosity, the fluorescence lifetime of the BOD-COOH derivative increased (figures 1D and 1F). To account for the variation of the lifetime as a function of the viscosity we used a model previously described by Vysniauskas *et al*.: ^33^

**Figure 1:**
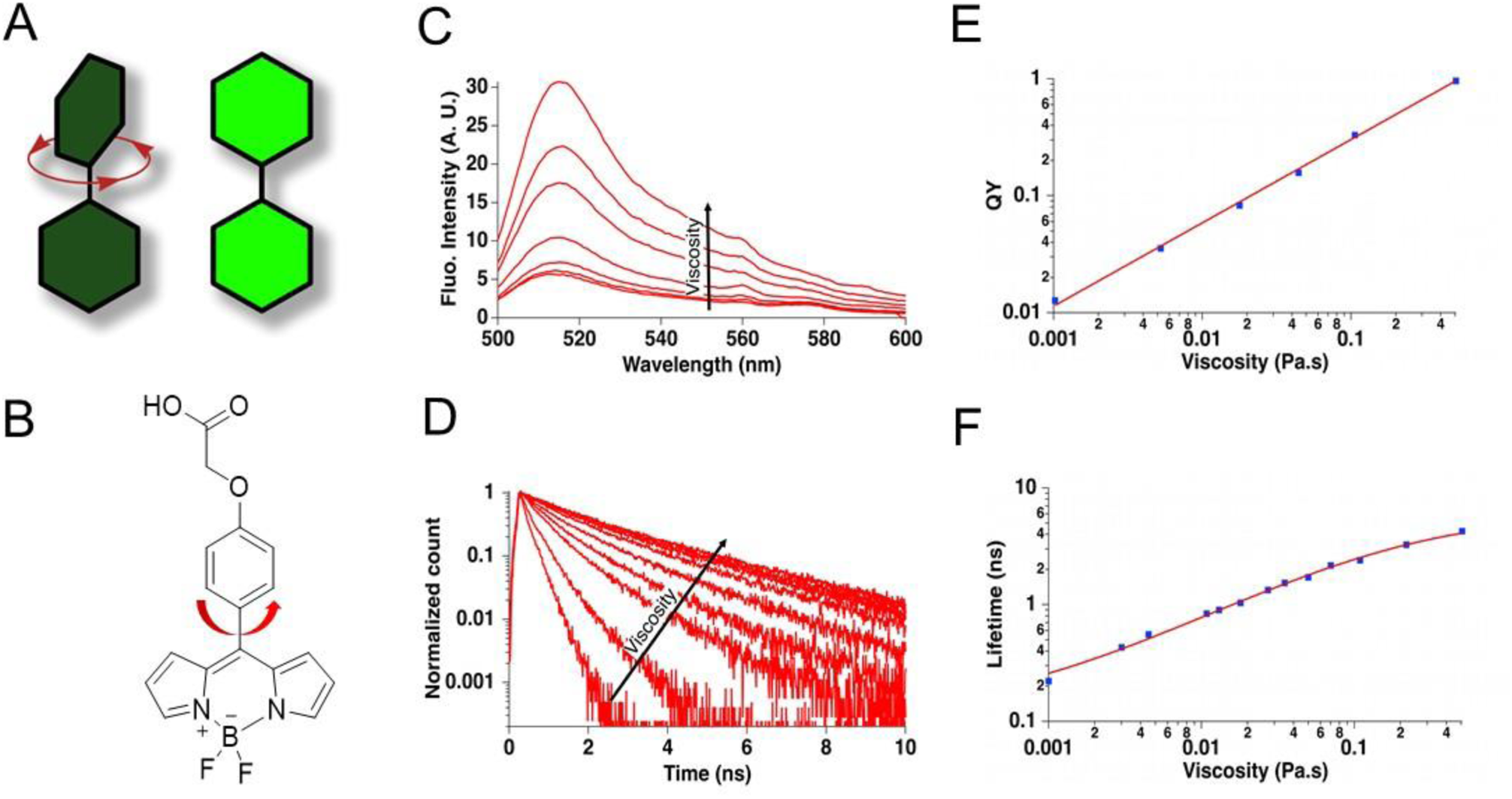
A). Rotational mechanism and structure of BODIPY-based molecular rotors, B) Structure of BOD-COOH C) The fluorescence emission spectra of BOD-COOH and D) Fluorescence decay curves of BOD-COOH in water/glycerol mixtures with increasing viscosities E-F) Foster-Hoffmann plots of fluorescence quantum yield and lifetime vs. viscosity (blue squares correspond to the data points and the continuous red lines to the fit obtained using equations 1 and 2).

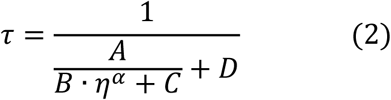

where A, B, C and D are unconstrained parameters that account for intrinsic lifetime measured at zero viscosity, the radiative lifetime of the dye (without non-radiative relaxation pathway) and the activation energy associated to the rotation of the phenyl moiety. *α* reflects the sensitivity of the molecular rotor to viscosity and was fixed to the value obtained from the quantum yield analysis.

To specifically target the BODIPY rotor to components of MLOs, we employed the HaloTag labeling strategy. A chloroalkane HaloTag ligand was directly conjugated to the fluorophore or linked via polyethylene glycol (PEG) spacers of four or twelve units (Figure 2A). The PEG linkers increase the distance between the BODIPY rotor and the binding site within the HaloTag binding pocket, thereby minimizing the influence of the protein environment on viscosity sensing. The time-resolved fluorescence decays of BOD-L, BOD-PEG4-L and BOD-PEG12-L were measured to retrieve their fluorescence lifetimes as a function of the viscosity of water/glycerol mixtures without (figure 2B) and in the presence of an excess of the purified HaloTag protein (figure 2C). As shown in Figure 2B, all three BODIPY derivatives retained viscosity sensitivity (α_*BOD*−*L*_ = 0.71 ± 0.04, α_*BOD*−*PEG*4−*L*_ = 0.64 ± 0.03, α_*BOD*−*PEG*12−*L*_ = 0.65 ± 0.04). Notably the viscosity sensitivity of BOD-L upon binding to HaloTag protein was lost (Figure 2C), indicating that the fluorophore remained inserted within the HaloTag binding pocket and was insufficiently exposed to the surrounding medium (α_*BOD*−*L*+*HaloTag*_ = 0.07 ± 0.01). In contrast, both BOD-PEG4-L and BOD-PEG12-L exhibited similar viscosity-dependent fluorescence responses, although their sensitivity was slightly reduced compared to the unbound fluorophores (α_*BOD*−*PEG*4−*L*+*HaloTag*_ = 0.29 ± 0.02, α_*BOD*−*PEG*12−*L*+*HaloTag*_ = 0.35 ± 0.05). This reduction suggests that rotational restriction imposed by the protein environment partially affects the fluorophore’s relaxation.

**Figure 2:**
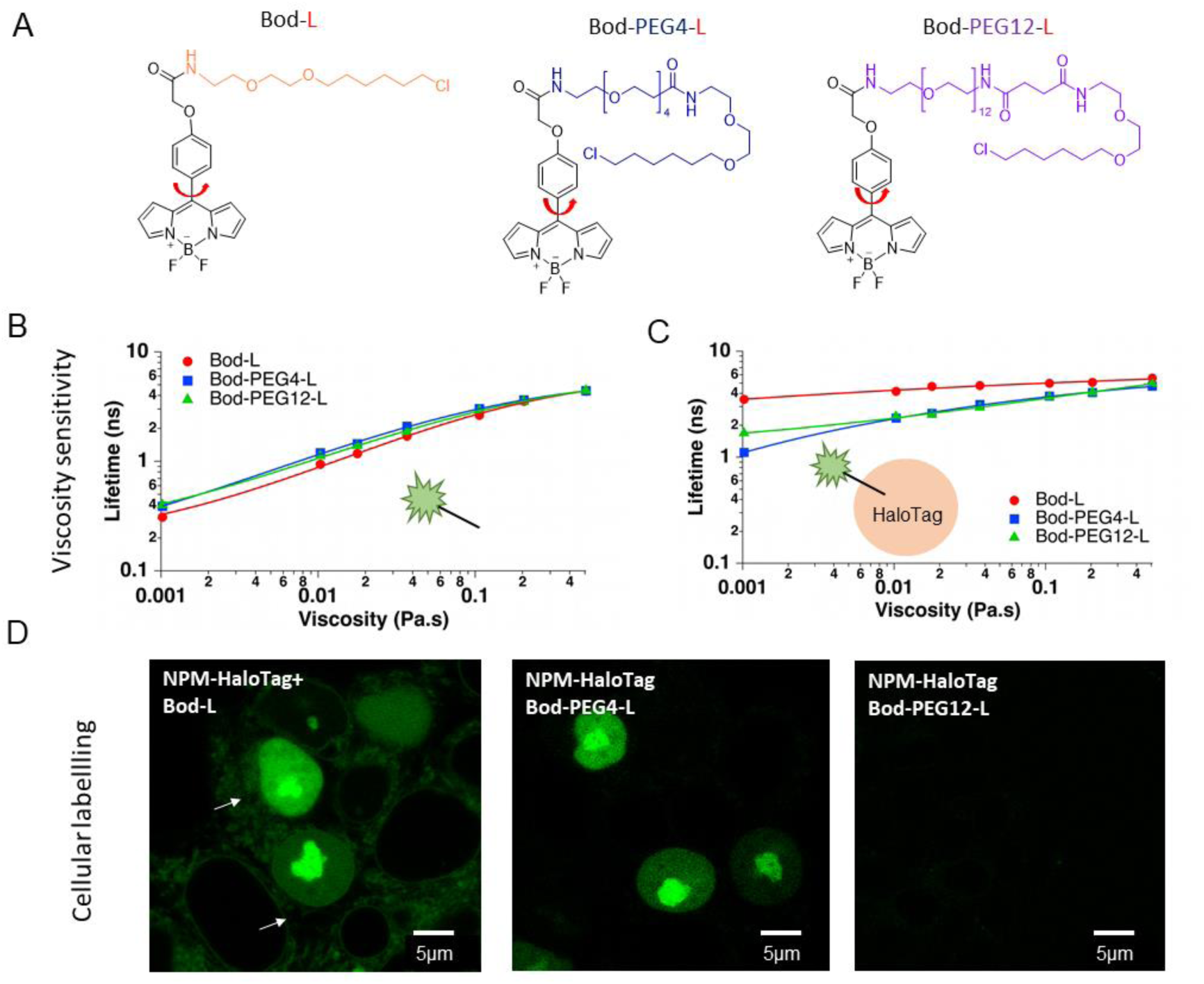
(A) Structures of BOD-L, BOD-PEG4-L and BOD-PEG12-L derivatives.( B) Fluorescence lifetimes measured for the BODs alone (50nM) (individual points correspond to data points and continuous line to the fit obtained using equation 2) and (C) in presence of HaloTag protein (0.5µM) (individual points correspond to data points and continuous line to the fit obtained using equation 2). For both conditions, the alpha values are reported in the main text.( D) HEK 293T cells expressing NPM-HaloTag labelled with BOD chloralkane derivatives. BOD-L labels specifically the cell nucleolus, however a slight cytoplasmic signal is also present. Cells labelled with BOD-PEG4-L show only specific probe binding to the NPM-HaloTag. For BOD-PEG12-L no fluorescence was detected in the cells.

We next evaluated the three BODIPY probes for their ability to label HaloTag fusion proteins in living cells. HEK293 cells expressing the nucleolar protein Nucleophosmin-1 fused to HaloTag (NPM-HaloTag) were labeled and imaged by confocal microscopy. BOD-L produced a strong fluorescence signal within the nucleolus but also exhibited substantial non-specific staining in the cytoplasm. In contrast, BOD-PEG4-L selectively labeled the nucleolus with minimal background fluorescence. No significant intracellular signal was detected for the BOD-PEG12-L derivative, which likely failed to penetrate the cells due to the increased hydrophilicity conferred by the long PEG12 linker. In light of these results, all cellular experiments in this study were performed with BOD-PEG4-L.

### BOD-PEG4-L senses the viscosity in various cellular compartments

To assess the sensitivity of the BOD-PEG4-L rotor to microviscosity within the cellular environment, different subcellular structures in U2OS cells were labeled by expressing specific HaloTag-fusion proteins: free cytoplasmic HaloTag protein, HaloTag-H2B in the cell nucleus, LifeAct–targeting HaloTag to actin filaments, and HaloTag–hFis1 in mitochondria. FLIM microscopy revealed distinct fluorescence lifetimes for each compartment. Based on the calibration curve reported in figure 2C, the average lifetime measured in the cytosol (τ_cytosol_ = 1.5 ns) revealed a low microviscosity (2.4 mPa.s).

Higher microviscosities were detected in the chromatin (τ_chromatin_ = 2.3 ns, 10 mPa.s) and the actin cytoskeleton (τ_actin_ = 2.5 ns, 14 mPa.s). The longest fluorescence lifetime was observed in hFis1 protein located in the mitochondrial outer membrane (τ_mitochondria_ = 3.5 ns, 71 mPa.s), indicating a more viscous environment. These values are in good agreement with previously reported viscosity estimations. ^34,35^

Altogether these data confirm that the sensitivity of BOD-PEG4-L probe is conserved in cells and that FLIM imaging reports on the microviscosity of the environment in the proximity of the labelled proteins in various cellular organelles.

### Mapping the microviscosity in different sub-compartments of the nucleolus

BOD-PEG4-L tool was next used to map the viscosity properties of the cell nucleolus. Nucleolus is a 1-2 µm sized MLO,^36^ it is implicated in the ribosomal biogenesis, cell cycle regulation and in the cellular stress response. Nucleolus is composed of three distinct compartments: the fibrillar centers (FC) – sites of rRNA transcription by RNA Pol 1, dense fibrillar component (DFC), where the rRNA are processed and the granular component (GC) being a site of the pre-ribosomal sub-units assembly.^37^

To monitor the microviscosity in the nucleolar sub-compartments, two nucleolar protein Nucleophosmin-1 (NPM) and Fibrillarin (Fib), were fused to HaloTag and expressed in HEK and U2OS cells. NPM localizes mainly in the GC, while Fib is preferentially located in the DFC. FLIM images were analyzed by decay fitting and also by phasor plot analysis. The phasor approach offers a fit-free, intuitive way to analyze lifetime data by converting the measured fluorescence decay in each pixel of the image, into a point (vector) in a 2D plot using a Fourier transform at the modulation frequency of the excitation source (see Mat and Methods section).^31,32^ In the resulting phasor plot the pixels are distributed on the semi-circle with the shortest lifetimes plotted on the right side of the plot and longer lifetimes closer the origin.^32^ Figure 3A represents Intensity images and corresponding phasor plots for NPM-HaloTang and Fib-HaloTag proteins expressed in HEK293. The phasor plots for each protein show elongated shape enabling to select the pixels of two populations with different lifetimes (Figure 4A, Phasor ROI and Phasor plot insets). When displaying corresponding pixels on the FLIM images, these two, populations represent NPM or Fib proteins located in the nucleoplasm (shorter lifetime) and in the nucleolus (longer lifetime).

**Figure 3:**
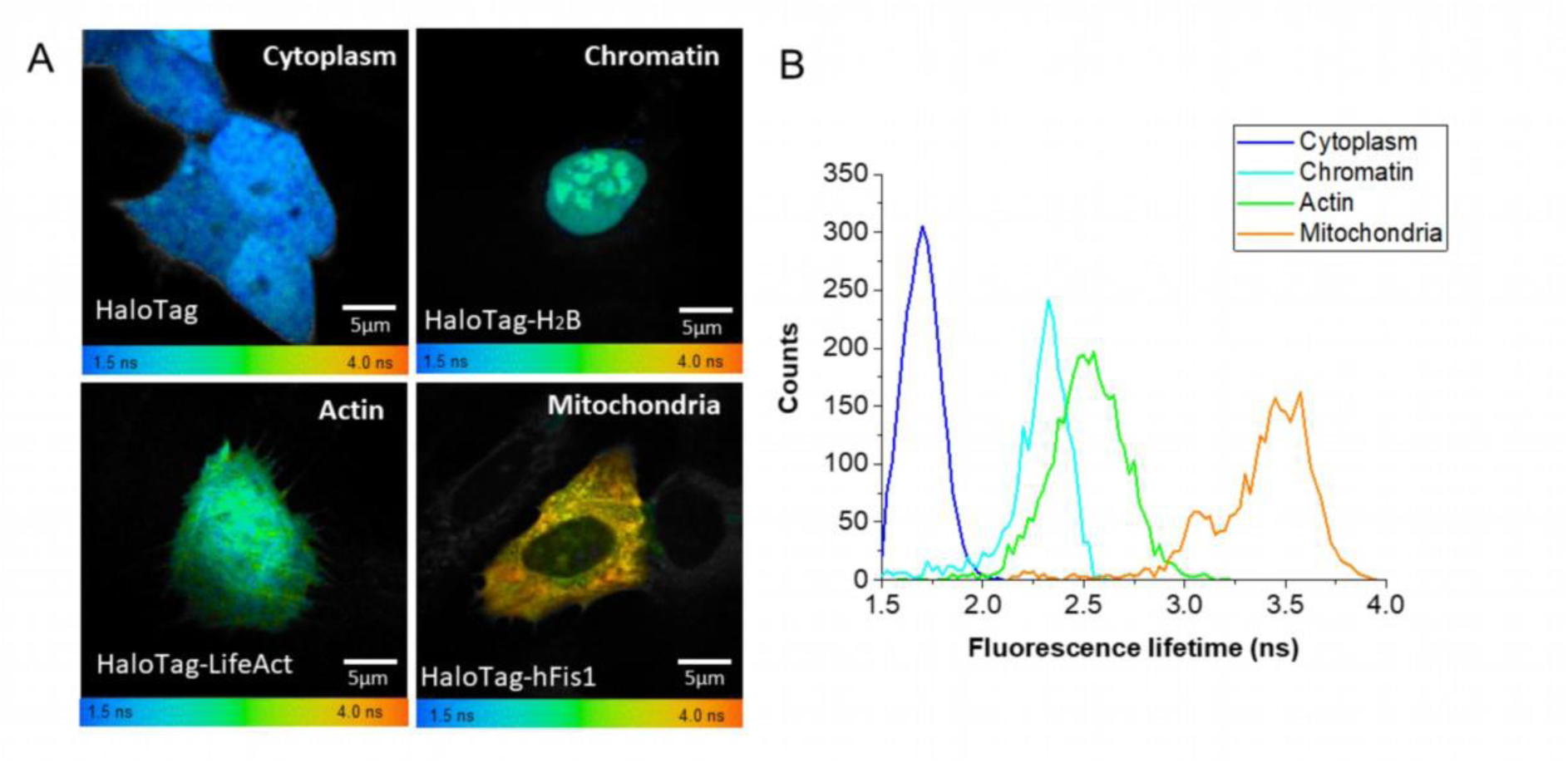
BOD-PEG4-L is sensitive to environment in various cellular compartments: (A) FLIM images of cells expressing free Halo Tag protein, HaloTag-H_2_B fusion (chromatin labelling), HaloTag-Life-Act to label the actin fibers and mitochondrial protein HaloTag-hFis1 (B). Corresponding Lifetime Distributions.

**Figure 4:**
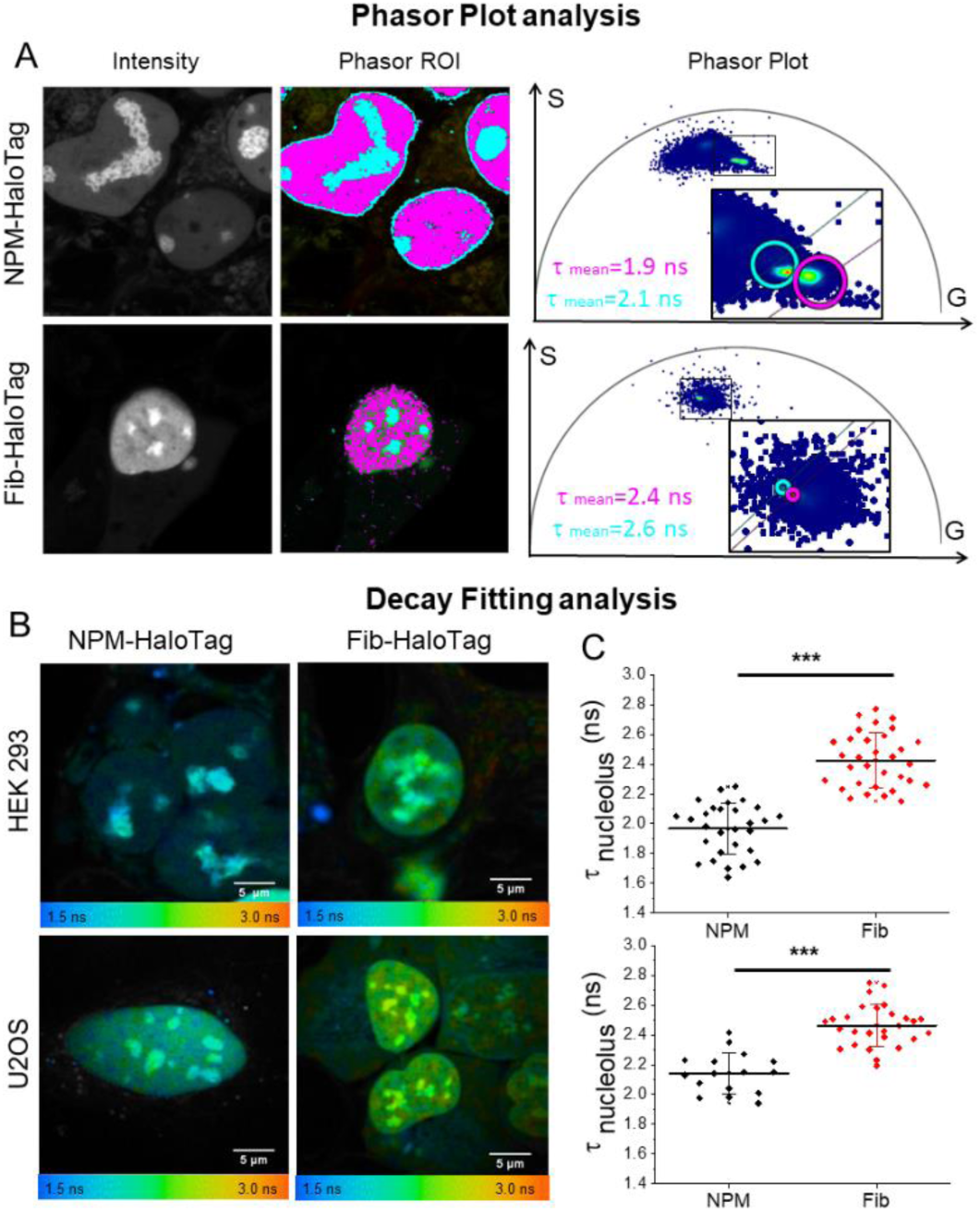
BOD-PEG4-L senses the microviscosity in nucleolar sub-compartments (A) Intensity images, phasor plot and phasor based ROI analysis of HEK293 cells expressing NPM-HaloTag or Fib-HaloTag labelled with BOD-PEG4-L. Phasor plot displays two populations with shorter lifetimes corresponding to the dye in the nucleoplasm and longer lifetimes for the dye present in the nucleolus. (B) FLIM images of HEK293 and U2OS cells expressing NPM-HaloTag and Fib-HaloTag. (C) Average lifetimes ± SD measured for 15-30 cells in each condition measured in 3 independent experiments, *** p<0.001.

BOD-PEG4-L bound to the nucleolar NPM-HaloTag showed a lifetime τ_NPM_=2.10± 0.06 ns, corresponding to 7 mPa.s. The lifetime of the probe bound to the fraction located in the nucleoplasm was τ_NPM_ = 1.88±0.06 ns corresponding to the viscosity of 4.7 mPa.s indicating different viscosities in both locations. Interestingly the environment within the DFC seems to be more viscous than GC, lifetimes measured for Fib-HaloTag are significantly higher compared to NPM (τ_Fib_ =2.60 ±0.04, 16 mPa.s in DFC and τ_Fib_ =2.42 ±0.05, 12 mPa.s in the nucleoplasm).

These trends were reproducible in two cell lines tested HEK293 and U2OS (see Table 1).

**Table 1:** Mean fluorescence lifetimes ± SD measured in nucleolar sub compartments and corresponding viscosity (η) values. Values based on 3 independent experiments with total 15-30 cells analyzed in each condition.

| Cell line | $\tau_{\text{GC}}$ (ns) | $\eta$ (mPa.s) | $\tau_{\text{GC}}$ (ns) +ActD | $\eta$ (mPa.s) | $\tau_{\text{DFC}}$ (ns) | $\eta$ (mPa.s) | $\tau_{\text{DFC}}$ (ns) +ActD | $\eta$ (mPa.s) |
| --- | --- | --- | --- | --- | --- | --- | --- | --- |
| HEK293 | 1.97±0.17 | 5.5 | 2.11±0.10 | 7 | 2.43±.18 | 12 | 2.62±0.25 | 16 |
| U2OS | 2.14±0.14 | 7.5 | 2.28±0.12 | 9.5 | 2.46±0.14 | 13 | 2.65±0.18 | 18 |

### Effect of rRNA transcription inhibition on nucleolar microviscosity

To test the sensitivity of BOD-PEG4-L probe to the viscosity modifications induced by the changes of the composition of the nucleolar sub-compartments, we performed the FLIM imaging of cells treated with Actinomycin D (ActD) an inhibitor of RNA polymerase I. In the treated nucleoli the DFCs fused together and progressively migrated to the nucleolar periphery, where they formed structures called “nucleolar caps”. Concomitantly, GC became smaller and spherical showing clearly its liquid like properties (Figure 5A). RNA imaging performed after 2h of ActD treatment showed no presence of RNA in the nucleolar caps; however, a residual RNA signal was still detectable in GCs (Figure 5B). Therefore, the DFC changed from a protein/RNA condensate to a purely protein-rich condensate, while the impact on the GC composition was less pronounced.

**Fig 5.**
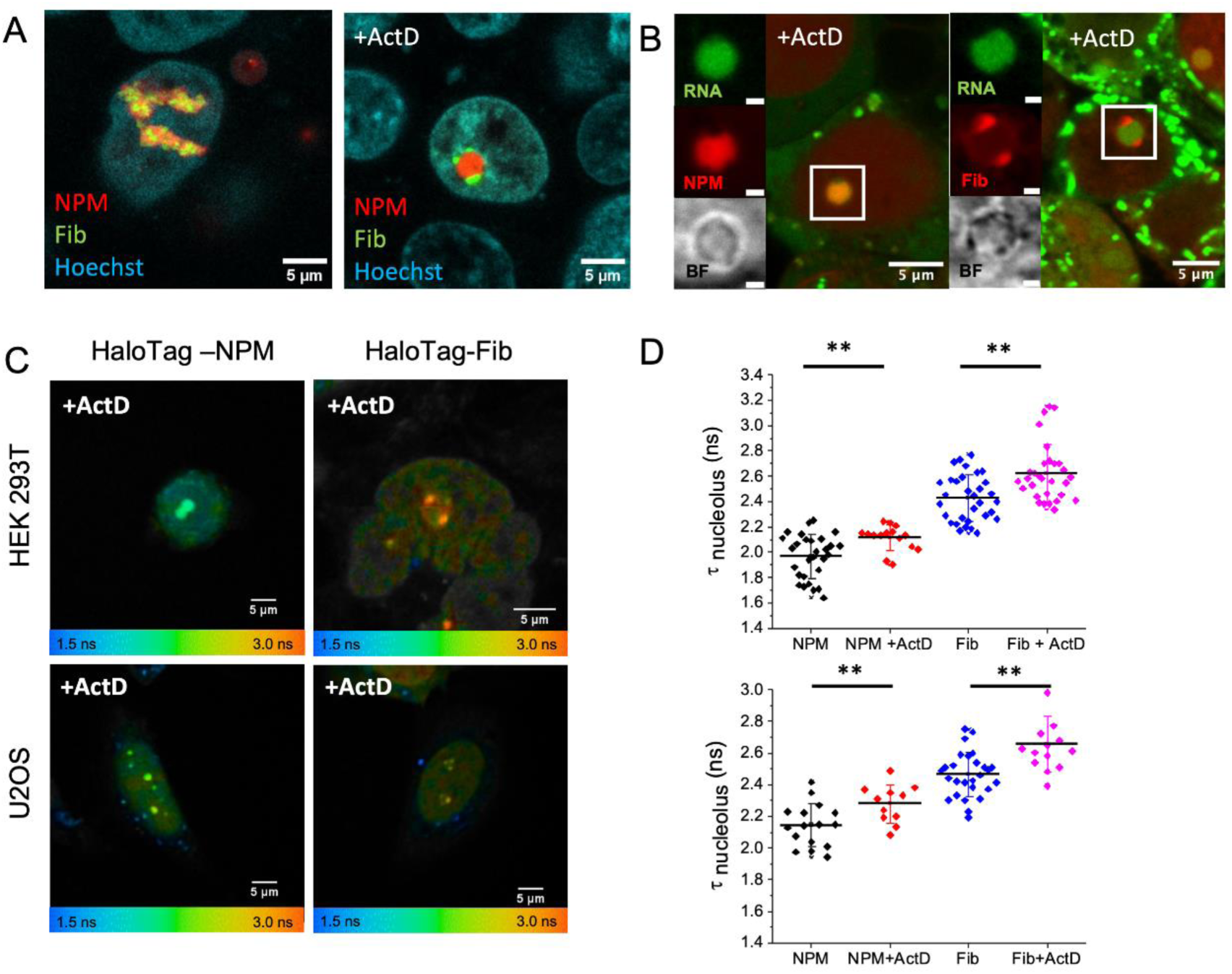
(A) Confocal images of HEK 293T cells expressing NPM-HaloTag (JF657-Halo), Fib-SNAP (JF552-SNAP). Act D treatment induces the separation of GC with DFC forming nucleolar caps on the surface. (B) Confocal images of ActD treated HEK293 cells expressing NPM-HaloTag (JF657-Halo) or Fib-HaloTag (JF657-Halo) co-stained with Pyrinon Y (1 μM, 15 min) (C) FLIM images HEK293 and U2OS cells expressing NPM-HaloTag or Fib-HaloTag labelled with BOD-PEG4-L. (D) Mean fluorescence lifetimes±SD. Values based on 3 independent experiments with total 15-30 cells analyzed in each condition, ** for p<0.01.

To monitor the microviscosity changes accompanying this nucleolar reorganization, HEK293T and U2OS cells expressing NPM-HaloTag or Fib-HaloTag labelled with BOD-PEG4-L were treated with ActD.

Two-photon fluorescence lifetime imaging (FLIM) confirmed a clear morphological change in the nucleolus upon drug treatment in both HEK293 AND U2OS cells. The lifetime measurements in ActD treated cells revealed an increase of BOD-PEG4-L fluorescence lifetime in both sub-compartments (Table 1 and Fig 5C and D). Fluorescence lifetime for NPM-HaloTag increased by 0.14 ns and 0.15 ns in HEK 293 and U2OS cells respectively, corresponding to a viscosity increase of around 2 mPa.s. Similarly, for the Fib-HaloTag an increase of 0.19 ns and 0.20 ns was measured which reflects the viscosity increase of 4 mPa.s.

This observation confirms that the reorganization of the condensate’s internal structure from RNA-containing to RNA-free domain has a direct impact on the internal environment sensed by the molecular rotor.

### Comparative Analysis of Nuclear and Cytoplasmic Condensates

Finally, we wanted to explore whether BOD-PEG4-L probe can sense the differences between cellular condensates. To this aim we performed a FLIM imaging of BOD-PEG4-L in stress granules (SG) in U2OS cells expressing Ras GTPase-activating protein-binding protein fused to HaloTag (HaloTag-G3BP1). G3BP1 is a major protein orchestrating the assembly of the SG.^38^ The latter are formed as a response of the cell to harsh environmental conditions such as oxidative and thermal stress or a presence of pathogens, when the cell induces a translational arrest in which cellular mRNA and RNA binding proteins concentrate in these liquid-like droplets.^39^

U2OS cells expressing HaloTag-G3PB1 were treated with sodium arsenite to induce the oxidative stress, labelled and imaged by FLIM. The average fluorescence lifetime measured in the stress granules is 1.45 ns (2.1 mPa.s, Figure 6B), corresponding to values measured for free HaloTag in the cell cytoplasm (Figure 3). The microviscosity in the SG was about 3.5 times lower as compared to the cell nucleolus. This result indicates that the RNA/protein network in the SG is sparse compared to the nucleolus and contains larger free spaces.

**Figure 6:**
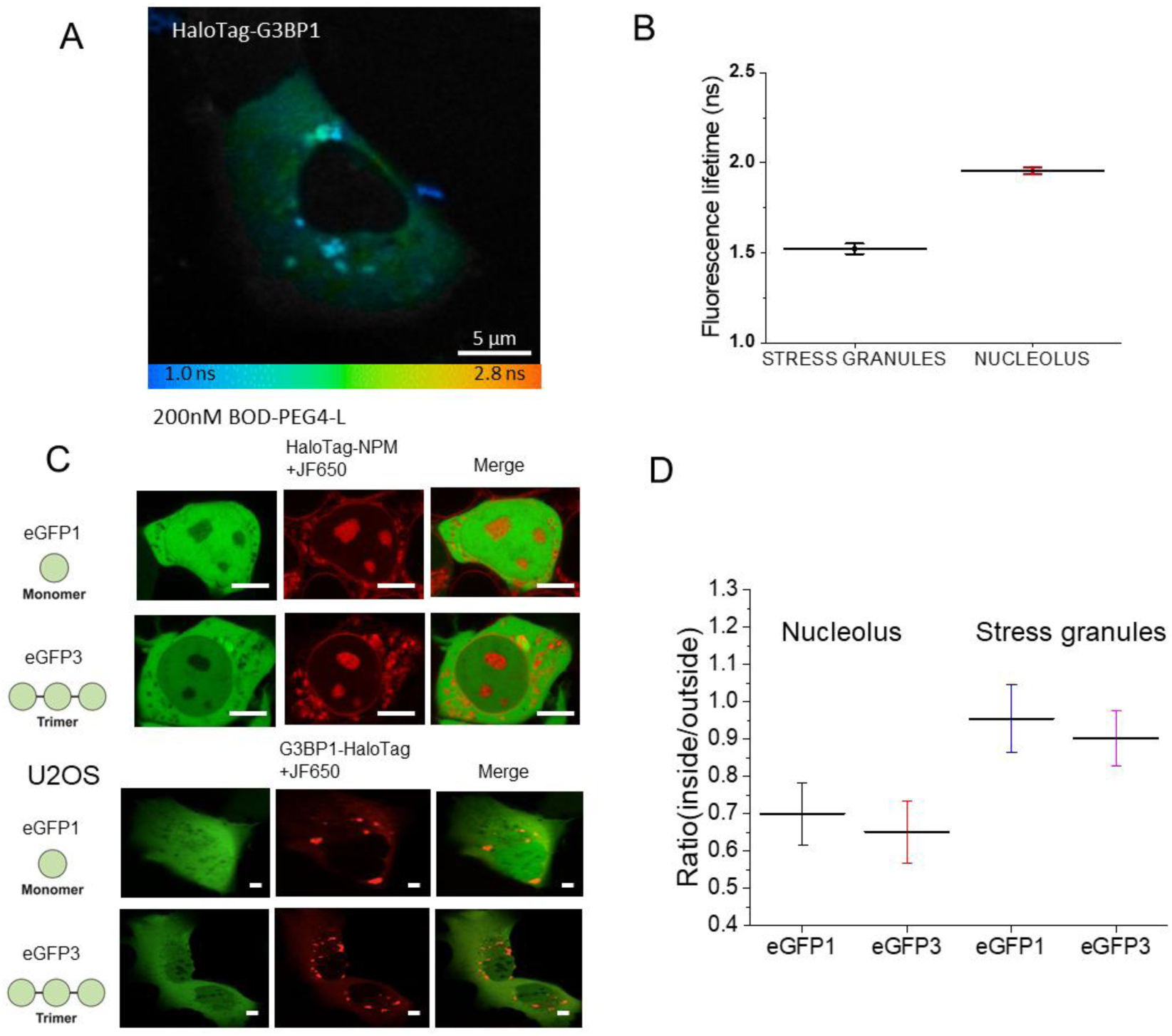
FLIM image of U2OS cells expressing HaloTag-G3BP1 labelled with BOD-PEG4-L. Cells were treated with Sodium Arsenite (0.5mM, 1h) to induce the formation of stress granules. (B) Mean fluorescence lifetimes±SD measured for NPM-HaloTag and HaloTag-G3BP1. Values based on 3 independent experiments with total 15-30 cells analyzed in each condition, (C) Confocal images of HEK293 cells expressing HaloTag-NPM labelled by JF650-Halo together with eGFP or eGFP3. U2OS cells expressing HaloTag-G3BP1 labelled by JF650-Halo together with eGFP or eGFP3. Scale bars: 5 µm (D) Fluorescence intensity ratios of eGFP signal inside vs. outside of both MLOs.

In order to verify this difference, U2OS cells were co-transfected with NPM-HaloTag or HaloTag-G3BP1 together with plasmids coding for eGFP or an eGFP trimer (eGFP3). The cells were imaged by confocal microscopy and the partition coefficient of eGFP and eGFP3 in both MLOs was quantified by measuring the ratio of eGFP fluorescence intensity outside vs. inside. Since eGFP monomer and trimer are not supposed to interact with the cellular structures, their diffusion within the MLOs in the cell is limited only by their size relative to protein/RNA meshwork density and permeability, hence the partition coefficient is a measure of permeability of the MLOs for each protein. eGFP and eGFP3 were found to be excluded from the nucleolus. The partition coefficient value of eGFP in the nucleolus was 0.7 and decreased to 0.6 for eGFP3. This observation indicates that the nucleolus is a relatively dense condensate since the monomeric eGFP with the hydrodynamic radius approaching 2.5 nm ^40^ is excluded. On the contrary, both eGFP and eGFP3 penetrated stress granules. The partition coefficient was close to unity for eGFP and decreased slightly to 0.9 for eGFP3 trimer, indicating the presence of much larger accessible free spaces within the stress granules.

These observations are in full agreement with the BOD-PEG4-L based FLIM imaging, indicating that compared to nucleolus, the stress granules are significantly less dense and less viscous MLOs.

## Discussion

In this study, we developed a BODIPY-based molecular rotor functionalized with a HaloTag ligand to enable direct monitoring of microviscosity in various cellular environments including membrane-less organelles (MLOs). To this end, we optimized the BODIPY rotor with a HaloTag ligand separated by a PEG4 linker. This design provided high sensitivity to environmental viscosity, cell permeability and efficient, specific labelling of cellular targets.

### Viscosity sensing using a HaloTag-Functionalized BODIPY Molecular Rotor

Using fluorescence lifetime imaging microscopy (FLIM), we first confirmed that the BOD-PEG4-L probe is sensitive to microviscosity across various cellular compartments. Our measurements indicated relatively low viscosity in the cell cytoplasm (η_cyt_ ≈ 5 mPa.s), in agreement with previous fluorescence correlation spectroscopy (FCS) studies.^41^ Interestingly, this value is lower than that reported by Kuimova and collaborators using a BODIPY rotor directly functionalized with a HaloTag ligand.^15^ The major difference between the two designs lies in the presence of the PEG4 linker in our probe, which spatially separates the rotor from the HaloTag protein. We therefore hypothesize that rotors lacking this linker may still be subject to partial steric or environmental constraints imposed by the protein, leading to an increased apparent viscosity. The viscosity measured by the probe targeted to H_2_B in the chromatin (η= 10 mPa.s) and actin fibers (η=14 mPa.s) was higher than that of the cytoplasm. FCS studies similarly reported reduced diffusion of monomeric eGFP in the nucleoplasm, by a factor of approximately 3.2 relative to aqueous solution,^35^ corresponding to an estimated viscosity of 3–4 mPa.s. Our measurements corroborate the notion that chromatin is embedded within a low-viscosity aqueous phase permissive to the diffusion of low-molecular-weight proteins. Fluorescence lifetimes recorded in mitochondria (τ ≈ 3.5 ns) were close to those reported by Chambers *et al.*^15^, although viscosity values inferred from calibration curves differed. These discrepancies likely arise from differences in the specific mitochondrial proteins targeted (matrix versus outer membrane), leading to distinct microenvironments.

### Quantification of Microviscosity in Nucleolar Subcompartments

After validating the viscosity sensitivity of the BOD-PEG4-L probe in diverse cellular contexts, we used this approach to quantify microviscosity within cellular MLOs. For both nucleolar proteins examined, we found that the nucleolus exhibits significantly higher viscosity than the surrounding nucleoplasm. Moreover, we observed clear differences between nucleolar subcompartments, the dense fibrillar component (DFC) displayed higher viscosity than the granular component (GC). Feric *et al*. showed by coalescence and FRAP measurements a liquid-like behaviour for GC and more viscoelastic behaviour for DFC in NPM and Fib reconstituted condensates and in cell nucleoli.^42^ Our results confirm the distinct viscosity properties among nucleolar subdomains, underscoring the high sensitivity of BOD-PEG4-L to its local environment. Overall viscosity values ranged from ∼5–8 mPa.s in the GC to ∼12–14 mPa.s in the DFC and appeared independent of the cell line examined. Our findings are consistent with nano scale techniques such as FCS measurements of GFPs diffusion in the nucleolus, which report diffusion coefficients reflecting a viscosity range of 4–26 mPa.s.^13,41,43^ However these values are considerably lower than viscosity measurements reported for *in vitro* reconstituted condensates ∼700 mPa s.^42^ Such differences stem from the distinct physical scales probed by each technique. Passive microrheology relies on tracking of fluorescent beads that are hundreds of nanometers in diameter. In these conditions, the beads diffusion is sensitive to the RNA–protein meshwork and long-range interactions that remain inaccessible to molecular-scale probes.

Finally, it should be noted that all measurements in this study were performed in asynchronous cell populations. Since the cell nucleolus disassembles before the cell division and reassembles at the end of the mitosis,^44,45^ it is highly probable that the nucleolar viscosity varies during the cell cycle, this question should be considered in future work.

### Transcriptional Stress and Nucleolar Remodelling

We further observed an increase in viscosity within both nucleolar compartments following inhibition of rRNA transcription with ActD. ActD treatment induces marked nucleolar reorganization: nucleolar volume decreases the GC adopts a rounded morphology and DFCs coalesce into nucleolar caps.^13^ These changes accompany the release of nucleolar proteins from their molecular complexes and alterations in their interactions with rRNA, leading to an increase in protein mobility and a partial relocalization to the nucleoplasm.^46^ In RNA-rich condensates, nucleic acids act as a polymer scaffold interconnecting proteins within a meshwork containing relatively large solvent-filled spaces. Reduction of RNA content leads to smaller, denser condensates, ^14^ increasing the probability of rotor–protein collisions. Consequently, viscosity measured in ActD-treated nucleoli is higher than in untreated, rRNA-rich nucleoli. Consistent with our observations, increases in nucleolar density following ActD treatment have been reported using optical diffraction tomography (ODT) and FCS. ^13,14^

### Microviscosity of Stress Granules Reveals a Low-Density RNA–Protein Network

Finally, we compared nucleolar viscosity with that of stress granules. Surprisingly, FLIM measurements indicated that SGs exhibit viscosities close to that of the cytoplasm (η_SG_ ≈ 2.1 mPa.s). This finding suggests that SGs possess a considerably more open RNA–protein meshwork than nucleolar compartments. This interpretation is supported by the high partitioning of monomeric and trimeric eGFP into SGs. Our results align with ODT analyses by Kim *et al.*, who showed that SGs provide minimal refractive-index contrast relative to the cytoplasm, indicating a low concentration of SG components.^14^ In line, single-molecule tracking of G3BP1 performed by Niewidok and co-workers, revealed the presence of immobile dense nanocores (100–200 nm), surrounded by a more fluid phase in which proteins remain highly mobile.^27^ Our results thus reinforce the view that SGs represent low-density cytoplasmic condensates with a large RNA meshwork.

In conclusion, this study introduces a microscopy-based approach for measuring microviscosity in living cells. The technique provides information complementary to existing methods for probing the material properties of MLOs. Given the key importance of rheological properties in the MLOs functions, the development of new analytical tools and experimental strategies is essential for advancing our understanding of condensate biophysics.

## Supporting information

Supplemental Methods contaning chemical synthesis and NMR spectra

## Acknowledgements

This work was funded by the French National Research Agency (ANR) FluoLLPS ANR-23-CE11-0016 and IdEx Recherche Exploratoire, Unistra (R22086MM). The authors would like to thank Imaging Center PIQ-QuESt (https://piq.unistra.fr/) and Plateforme d’imagerie “In Vitro”, members of the national infrastructure France-BioImaging supported by the French National Research Agency (ANR-24-INBS-0005 FBI BIOGEN).

## Notes

### Competing Interest Statement

The authors have declared no competing interest.

