## Supplemental Methods contaning chemical synthesis and NMR spectra for "Fluorescence based microviscosity mapping in membraneless organelles"

1. **Synthesis and NMR spectroscopy of Bodipy derivatives:**

**BODIPY-COOH:**

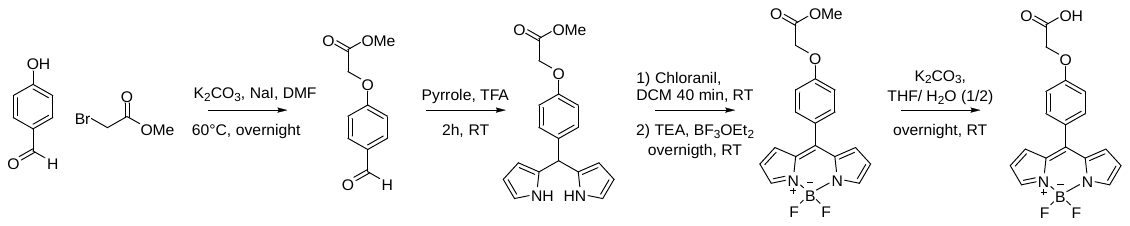

BODIPY-COOH was synthesized according to our previous reported protocol.^1^

**BODIPY-COOH:** ^1^H NMR (400 MHz, CD_3_OD-CDCl_3_ _1_-_1_) δ 7.82 (s, 2H), 7.43 (d, *J* = 8.8 Hz, 2H), 7.01 (d, *J* = 8.7 Hz, 2H), 6.87 (d, *J* = 4.2 Hz, 2H), 6.48 (dd, *J* = 4.3, 2.0 Hz, 2H), 4.66 (s, 2H), 3.31 (s, 1H).

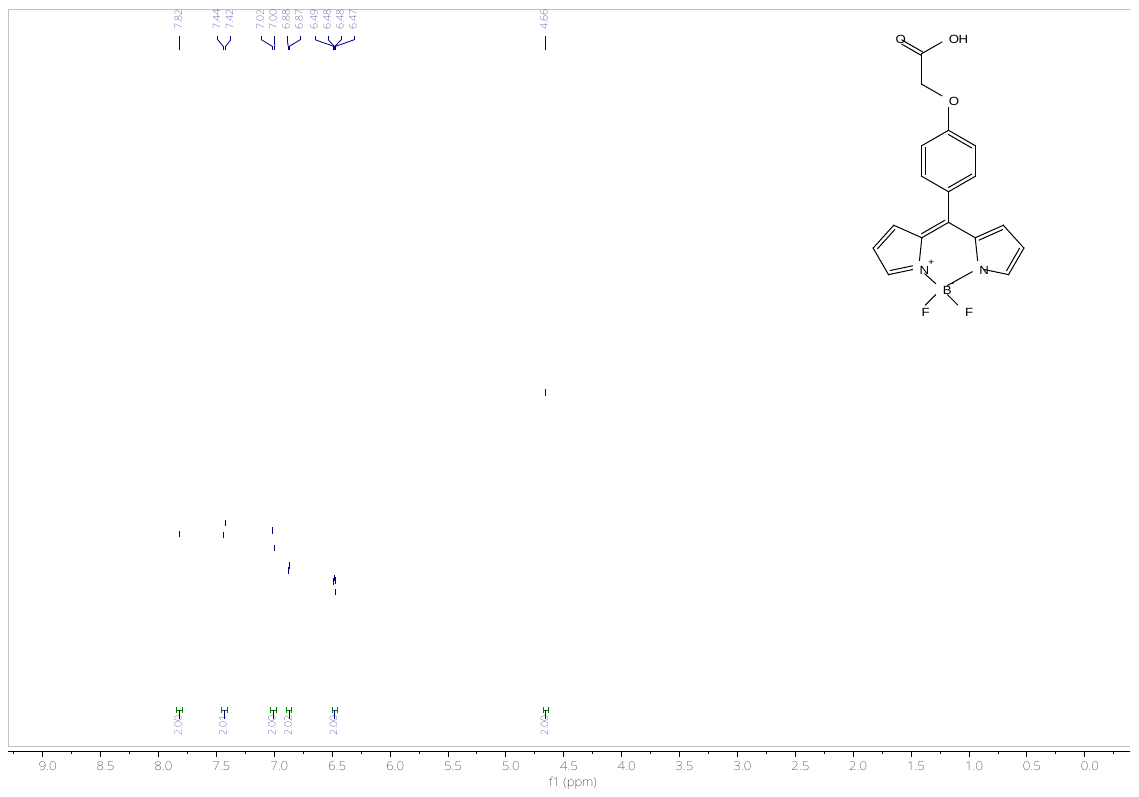

^1^H NMR **BODIPY-COOH**

**BOD-L:**

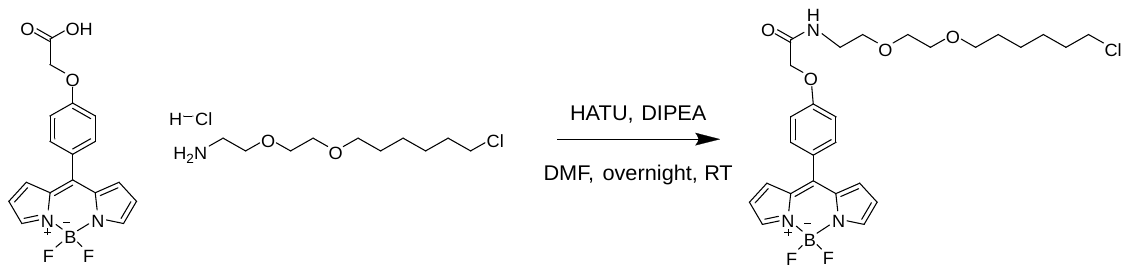

**BOD-L:** BODIPY-COOH (39.80 mg, 0.116 mmol, 1 equiv.), DIPEA (61 µL, 0.35 mmol, 3 equiv.) and 2-(2-((6-chlorohexyl)oxy)ethoxy)ethan-1-amine hydrochloride (30.3mg, 0.12 mmol, 1 equiv.) were dissolved in dry DMF (1 mL). HATU (49 mg, 0.13 mmol, 1.1 equiv.) was added to the reaction mixture. The reaction mixture was stirred overnight at room temperature. The mixture was diluted with DCM and washed with water and brine, dried over MgSO_4_, filtered, and the solvent was removed in vacuo. The residue was purified by flash column chromatography (0-10% MeOH/DCM) to give BOD-L as an orange oil (6 mg, 0.01 mmol, 9.4 %).

**^1^H NMR (400 MHz, CDCl3)** δ 7.93 (s, 2H), 7.57 (d, J = 8.7 Hz, 2H), 7.08 (d, J = 8.7 Hz, 2H), 6.94 (d, J = 4.2 Hz, 2H), 6.58 – 6.51 (m, 2H), 4.60 (s, 2H), 3.64 – 3.60 (m, 5H), 3.59 – 3.56 (m, 2H), 3.53 – 3.44 (m, 4H), 1.75 (dd, J = 8.0, 6.6 Hz, 2H), 1.65 – 1.55 (m, 4H), 1.46 – 1.33 (m, 4H).

**^13^C NMR (126 MHz, CDCl3)** δ 167.46, 159.49, 146.68, 143.88, 134.84, 132.47, 131.30, 127.69, 118.51, 114.84, 71.31, 70.40, 70.05, 69.70, 67.49, 44.99, 38.91, 32.49, 29.45, 26.66, 25.41.

**HRMS m/z:** [M-F]^+^ calculated for C_27_H_33_BClFN_3_O_4_: 528.2237, found 528.2247

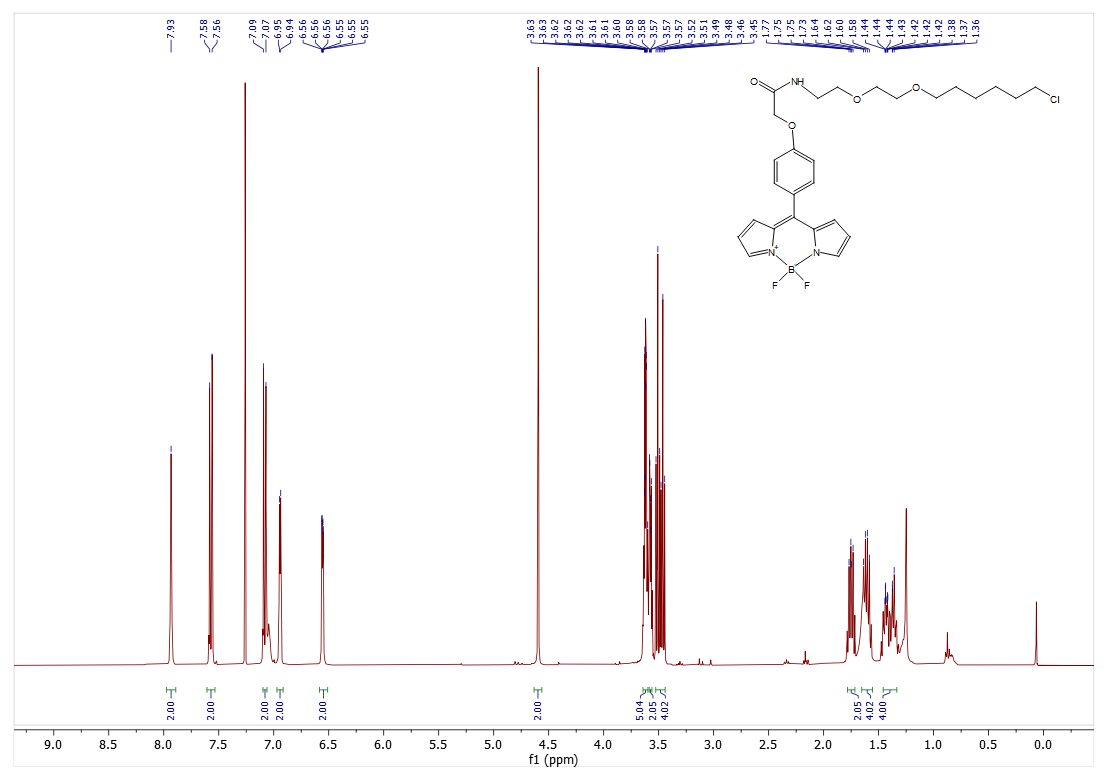

^1^H NMR of **BOD -L**

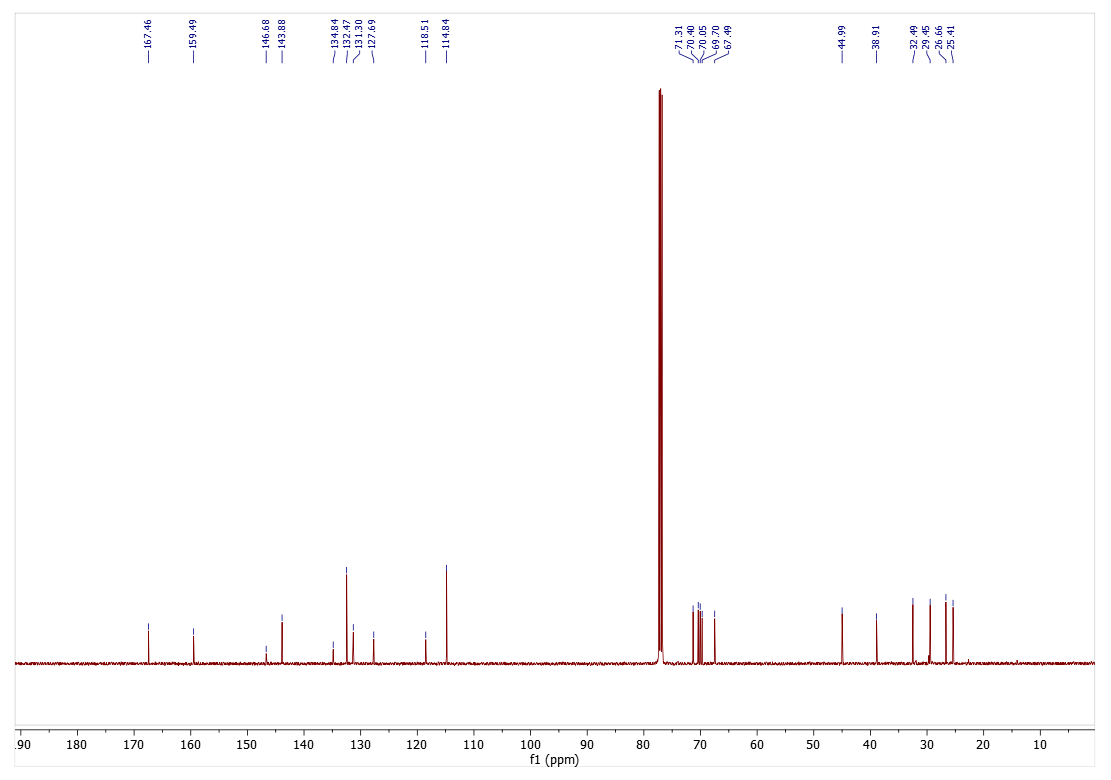

^13^C NMR of **BOD -L**

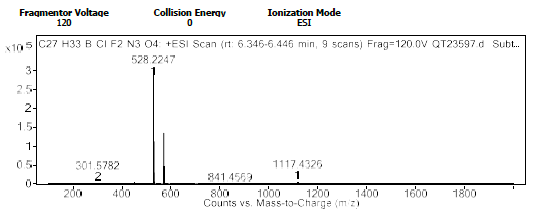
 HRMS of **BOD -L**

***1.1 General Procedure for Halo-PEG synthesis:***

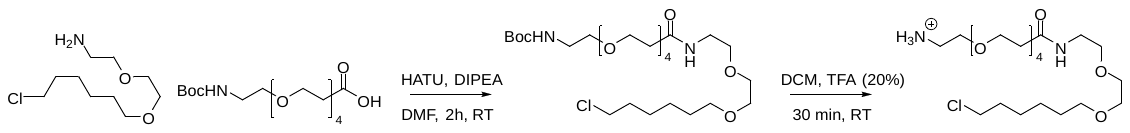

**Boc-Halo-PEG-4**^2^: To a solution of 2,2-dimethyl-4-oxo-3,8,11,14,17-pentaoxa-5-azaicosan-20-oic acid (130 mg, 581 µmol, 1.0 equiv.) and 2-(2-((6-chlorohexyl)oxy)ethoxy)ethan-1-amine.HCl (212 mg, 581 µmol 1 mmol, 1 equiv.) in dry DMF (2.5 mL) was added DIPEA (305 µL, 1.7 mmol, 3 equiv.) and the mixture was stirred at room temperature for 5 minutes. HATU (287 mg, 0.7 mmol, 1.3 equiv.) was then added and the mixture was stirred at room temperature for 2 hours. The reaction mixture was concentrated and diluted with DCM and washed with water and brine, dried over MgSO4, filtered, and the solvent was removed in vacuo. The residue was purified by flash column chromatography (0-10% MeOH/DCM) to yield tert-butyl (28-chloro-15-oxo3,6,9,12,19,22-hexaoxa-16-azaoctacosyl)carbamate as a colourless oil (180 mg, 0.56 mmol, 96 %).

**^1^H NMR (400 MHz, MeOD)** δ 3.74 (t, *J* = 6.2 Hz, 2H), 3.67 – 3.54 (m, 20H), 3.51 (q, *J* = 6.2 Hz, 4H), 3.39 (td, *J* = 5.5, 4.1 Hz, 2H), 3.26 – 3.19 (m, 2H), 2.47 (t, *J* = 6.2 Hz, 2H), 1.83 – 1.74 (m, 2H), 1.66 – 1.57 (m, 2H), 1.54 – 1.36 (m, 13H).

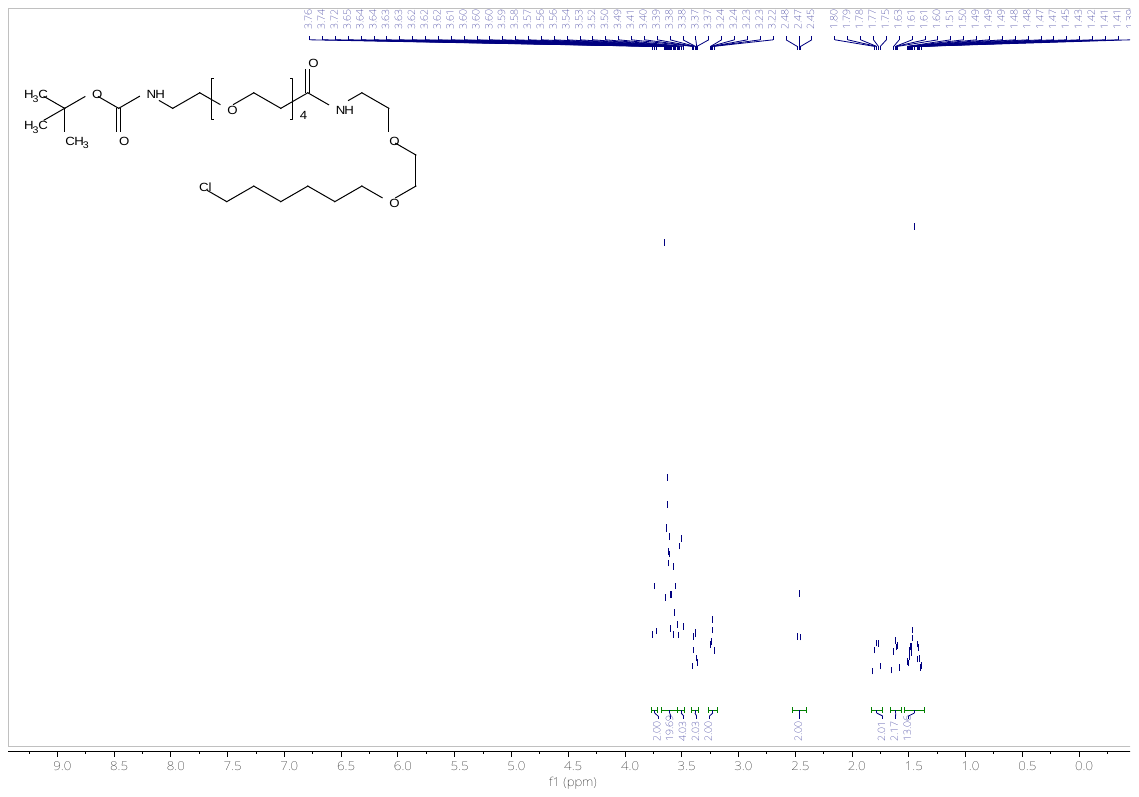

^1^H NMR **Boc-Halo-PEG-4**

**Amino-Halo-PEG-4**^2^: To a solution of tert-butyl (28-chloro-15-oxo-3,6,9,12,19,22-hexaoxa-16- azaoctacosyl)carbamate (75 mg, 0.13 mmol) in DCM (1 mL), TFA (200 µL) was added, and the mixture was stirred at room temperature for 30 minutes. The solvent was removed in vacuo to give 1-amino-N-(2-(2-((6-chlorohexyl)oxy)ethoxy)ethyl)- 3,6,9,12-tetraoxapentadecan-15-amide as a colourless oil (53 mg, 0.11 mmol, 85 %). The product was use without purification in the next step.

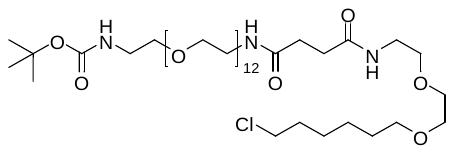

**Boc-Halo-PEG-12:** was obtain as a colourless oil (66 mg, 0.08 mmol, 100 %).

**^1^H NMR (500 MHz, CDCl3)** δ 6.58 (s, 1H), 6.45 (s, 1H), 5.07 (s, 1H), 3.67 – 3.50 (m, 56H), 3.47 – 3.39 (m, 6H), 3.32– 3.25 (m, 2H), 2.50 (s, 4H), 1.76 (p, J = 7.0 Hz, 2H), 1.59 (p, J = 7.0 Hz, 2H), 1.48 – 1.39 (m, 11H), 1.39 – 1.33 (m, 2H).

**^13^C NMR (500 MHz, CDCl_3_)** δ 172.35, 172.29, 156.14, 150.68, 79.24, 71.38, 70.63, 70.61, 70.41, 70.35, 70.14, 69.90, 69.86, 45.14, 40.48, 39.43, 39.39, 32.62, 31.79, 31.69, 29.54, 28.54, 26.78, 25.51.

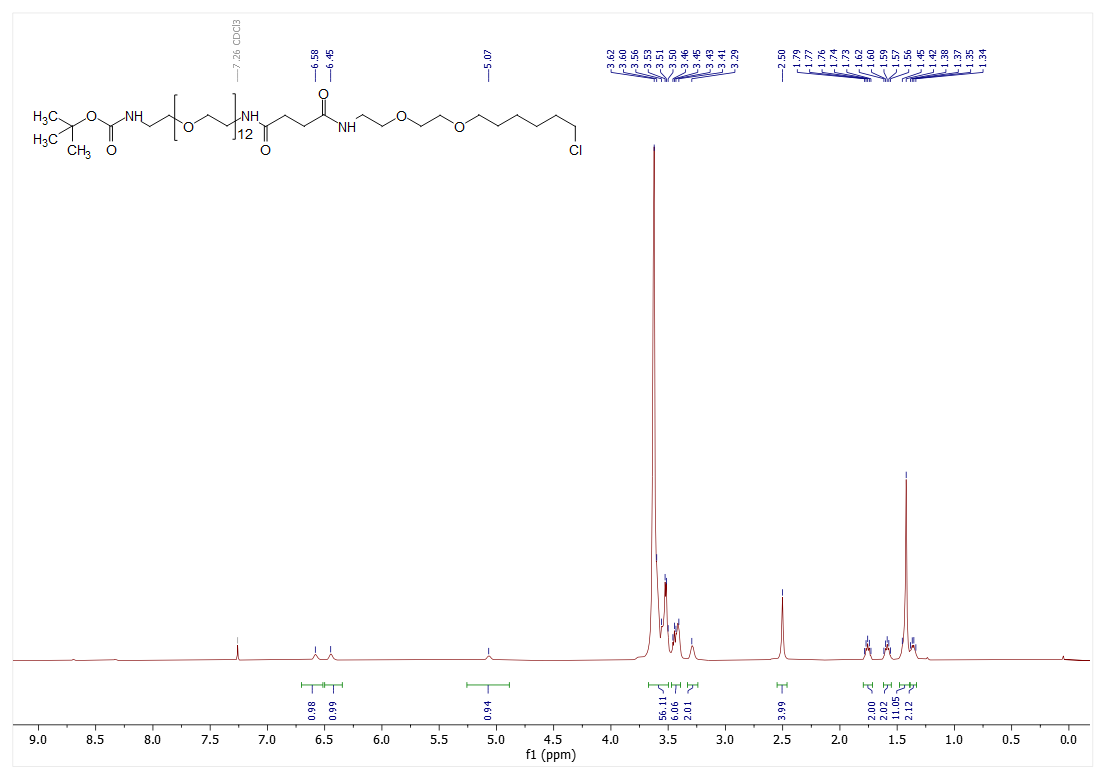

^1^H NMR **Boc-Halo-PEG-12**

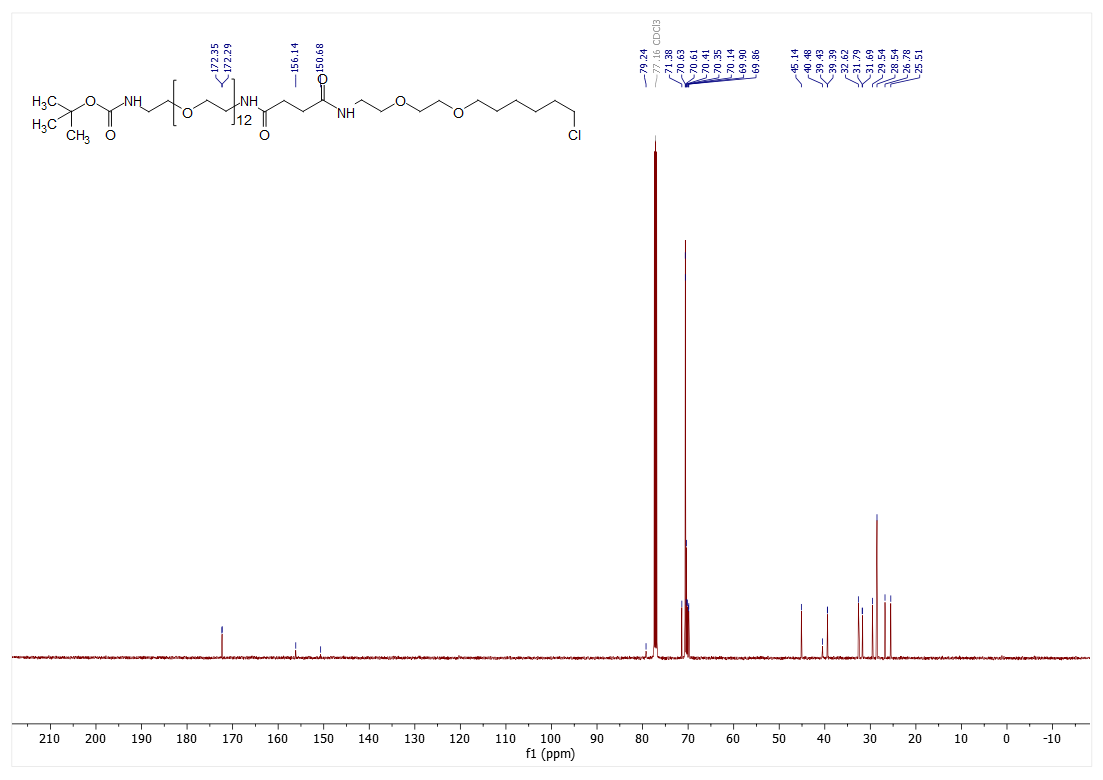

^13^C NMR **Boc-Halo-PEG-12**

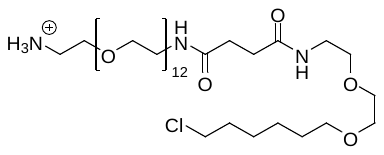

**Amino-Halo-PEG-12:** was obtain as a colourless oil (59 mg, 0.07 mmol, 100 %). The product was use without purification in the next step.

***1.2 Typical Procedure for BODIPY-COOH coupling with amino-Halo-PEG:***

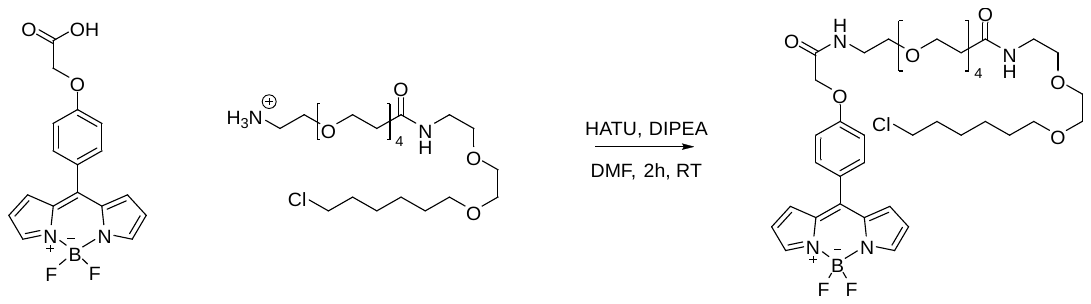

**BOD-PEG4-L:** BODIPY-COOH (10.7 mg, 0.03 mmol, 1 equiv.), DIPEA (17 µL, 0.09 mmol, 3 equiv.) and 1-amino-N-(2-(2-((6-chlorohexyl)oxy)ethoxy)ethyl)- 3,6,9,12-tetraoxapentadecan-15-amide (24 mg, 0.04 mmol, 1.3 equiv.) were dissolved in dry DMF (1.5 mL). HATU (15.5 mg, 0.04 mmol, 1.0 equiv.) was added to the reaction mixture. The reaction mixture was stirred at room temperature for 2 hours. The mixture was diluted with DCM and washed with water and brine, dried over MgSO_4_, filtered, and the solvent was removed in vacuo. The residue was purified by flash column chromatography (0-10% MeOH/DCM) to give **BOD-PEG4-L** as an orange oil (18.8 mg, 0.02 mmol, 75 %).

**^1^H NMR (400 MHz, CDCl_3_)** δ 7.81 (s, 2H), 7.48 – 7.41 (m, 2H), 7.01 – 6.96 (m, 2H), 6.83 (d, *J* = 4.2 Hz, 2H), 6.44 (dd, *J* = 4.3, 1.8 Hz, 2H), 4.48 (s, 2H), 3.61 (t, *J* = 5.9 Hz, 2H), 3.55 – 3.49 (m, 15H), 3.48 (p, *J* = 2.9 Hz, 4H), 3.45 – 3.41 (m, 5H), 3.40 (d, *J* = 6.7 Hz, 2H), 3.32 (td, *J* = 6.0, 2.7 Hz, 4H), 2.34 (t, *J* = 6.0 Hz, 2H), 1.68 – 1.61 (m, 2H), 1.50 – 1.44 (m, 2H), 1.35 – 1.30 (m, 2H), 1.27 – 1.20 (m, 2H).

**^13^C NMR (101 MHz, CDCl_3_)** δ 171.42, 167.72, 159.59, 146.77, 143.81, 134.83, 132.47, 131.34, 127.59, 118.46, 114.88, 71.25, 70.47, 70.44, 70.39, 70.31, 70.27, 70.25, 70.18, 70.02, 69.90, 69.74, 67.43, 67.26, 45.03, 39.18, 38.95, 36.87, 32.51, 29.45, 26.67, 25.41.

**HRMS m/z:** [M+Na]^+^ calculated for C_38_H_54_BClF_2_N_4_NaO_9_: 817.3538, found 817.3527.

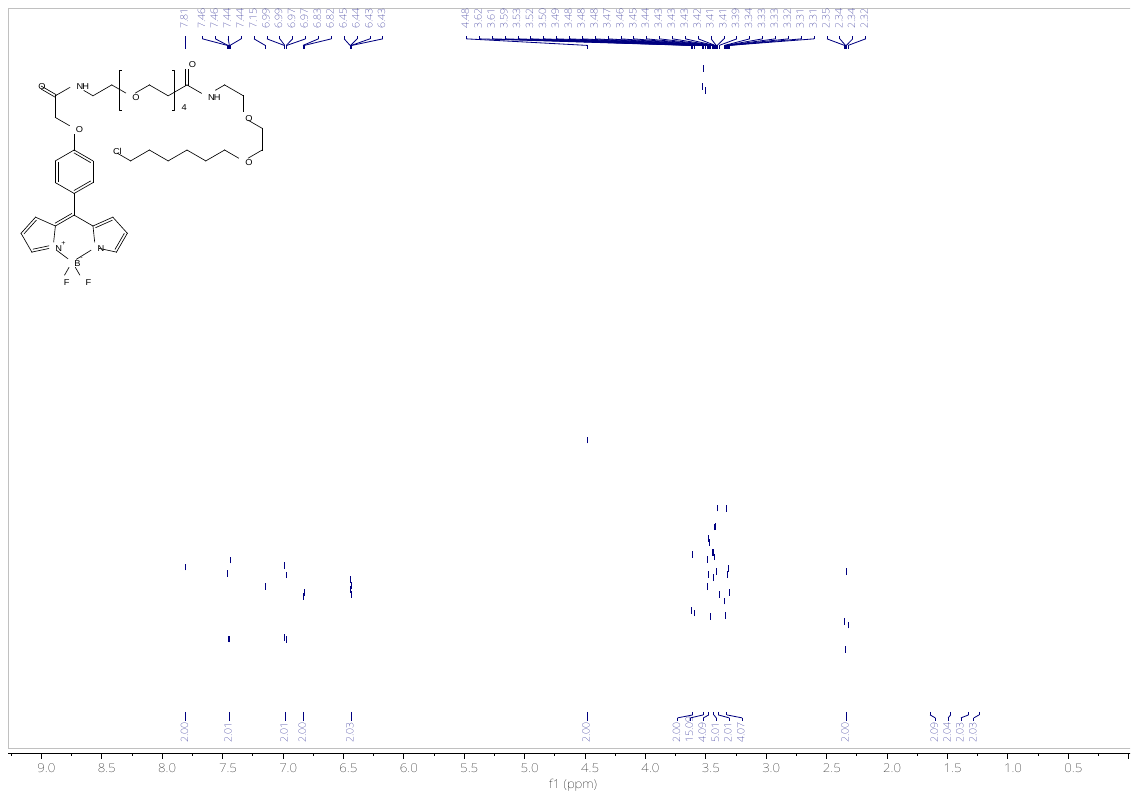

^1^H NMR of **BOD-PEG4-L**

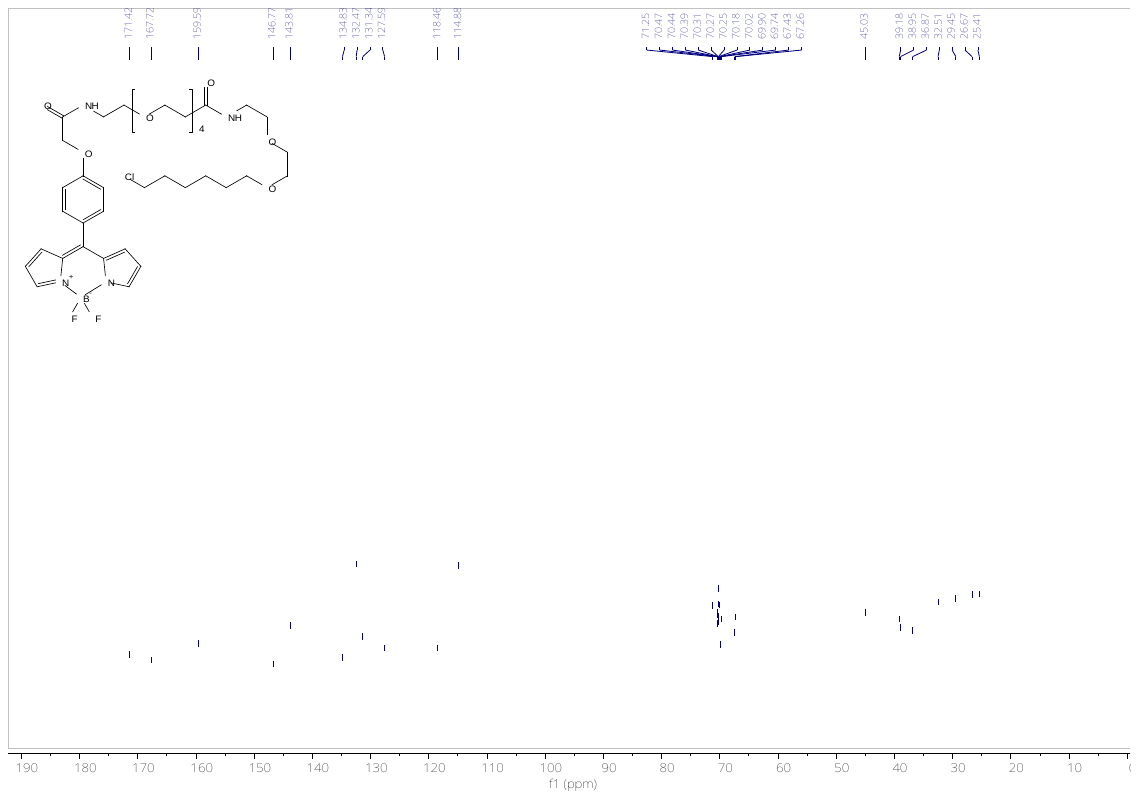
^13^C NMR of **BOD-PEG4-L**

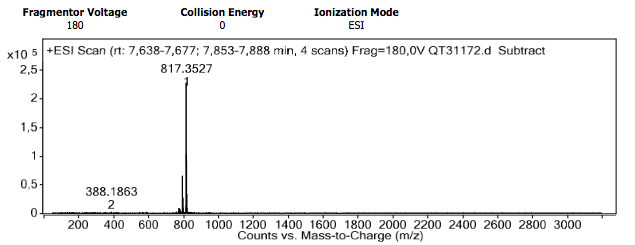

HRMS of **BOD-PEG4-L**

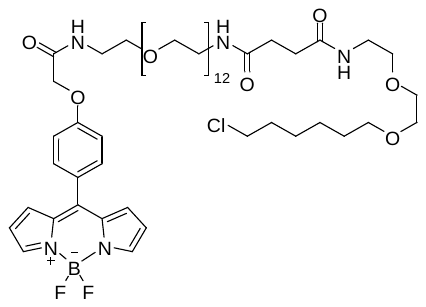

**BOD-PEG12-L:** was obtain as an orange oil (12mg, 0.09 mmol, 45%).

**^1^H NMR (400 MHz, CDCl_3_)** δ (ppm) 7.93 (s, 2H), 7.57 (d, *J* = 7.9 Hz, 2H), 7.09 (d, *J* = 8.4 Hz, 2H), 6.95 (d, *J* = 4.1 Hz, 2H), 6.56 (dd, *J* = 4.2, 1.8 Hz, 2H), 6.45 (s, 1H), 6.35 (s, 1H), 4.60 (s, 2H), 3.69 – 3.61 (m, 57H), 3.54 (d, *J* = 6.5 Hz, 6H), 3.45 (d, *J* = 7.0 Hz, 4H), 2.51 (s, 4H), 1.96 (s, 2H), 1.82 – 1.73 (m, 3H), 1.60 (t, *J* = 7.2 Hz, 3H), 1.50 – 1.33 (m, 6H).

**^13^C NMR (101 MHz, CDCl_3_)** δ 172.09, 167.48, 159.56, 143.85, 134.84, 132.49, 131.35, 127.64, 118.48, 114.89, 71.30, 70.61, 70.60, 70.38, 70.34, 70.07, 69.82, 69.72, 67.51, 45.05, 39.35, 39.29, 38.96, 32.54, 31.71, 31.63, 29.70, 29.47, 26.69, 25.43. *Some quaternary carbon signals were not observed in the ^13^C NMR spectrum despite extended acquisition times, likely due to long relaxation times and low sensitivity.*

**HRMS m/z:** [M+Na]^+^ calculated for C_57_H_91_BClF_2_N_5_NaO_18_: 1240.6006, found 1240.5980.

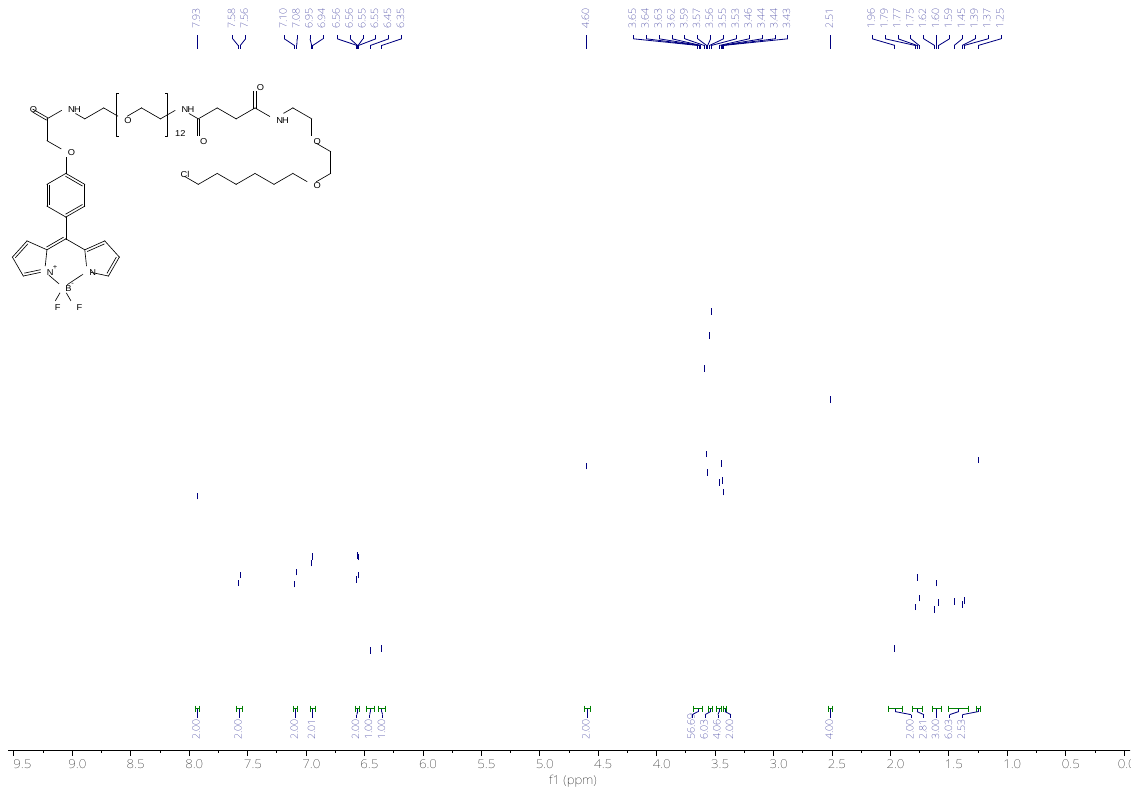

^1^H NMR of **BOD-PEG12-L**

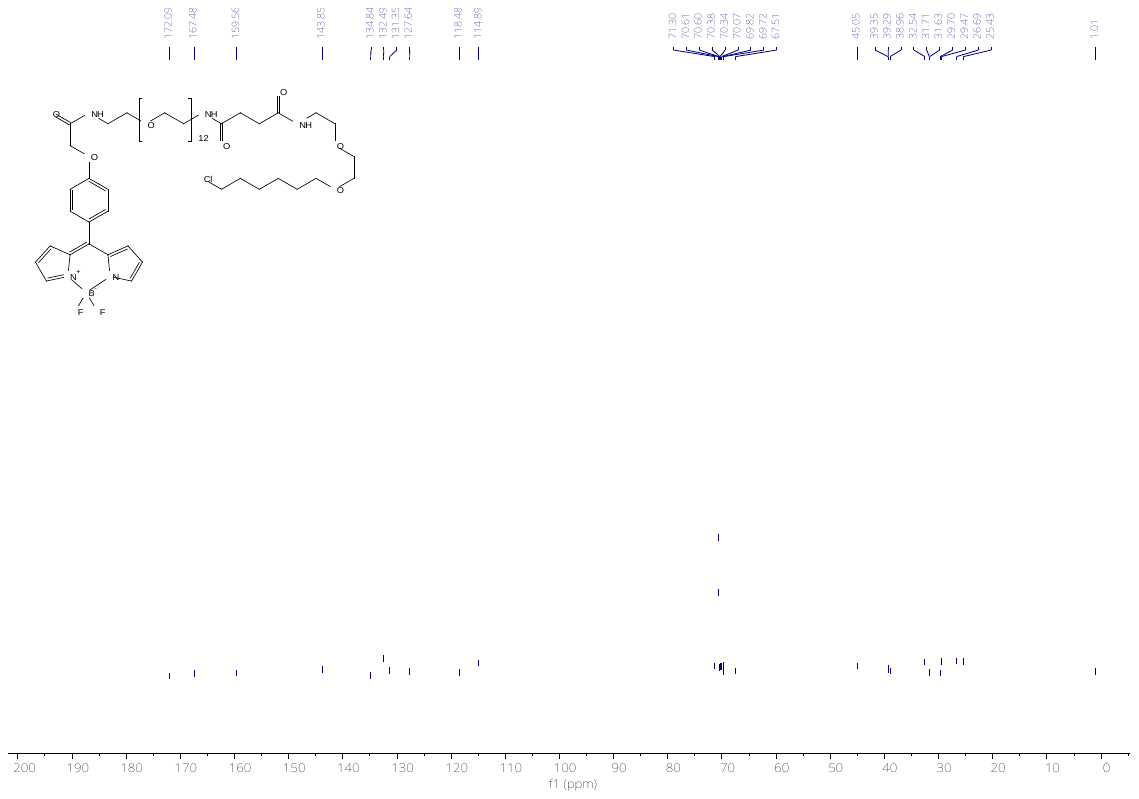

^13^C NMR of **BOD-PEG12-L**

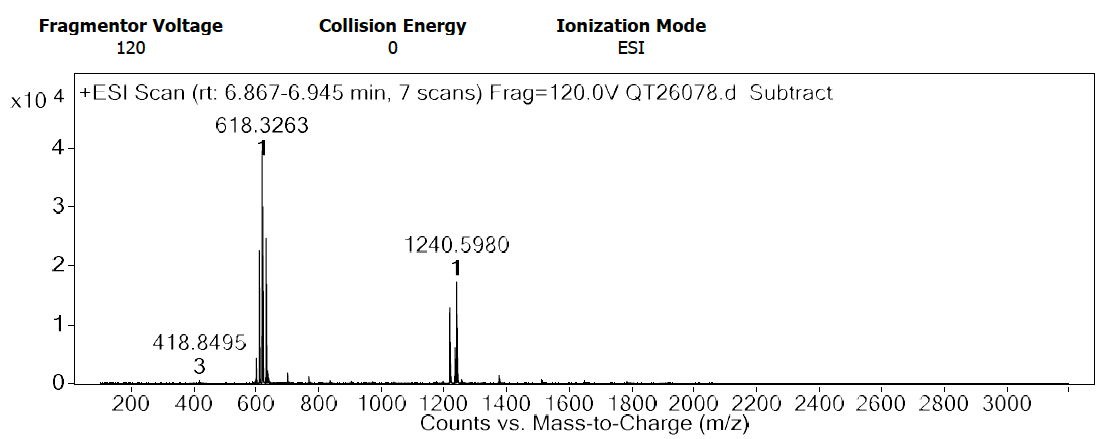

HRMS of **BOD-PEG12-L**

(1)	Ashokkumar, P.; Ashoka, A. H.; Collot, M.; Das, A. A FLUOROGENIC BODIPY MOLECULAR ROTOR AS APOPTOSIS MARKER. 22.

(2)	Simpson, M. M.; Lam, C. C.; Goodman, J. M.; Balasubramanian, S. Selective Functionalisation of 5-Methylcytosine by Organic Photoredox Catalysis. *Angew. Chem. Int. Ed.* **2023**, *62* (26), e202304756. https://doi.org/10.1002/anie.202304756.
